# Phasing Diploid Genome Assembly Graphs with Single-Cell Strand Sequencing

**DOI:** 10.1101/2024.02.15.580432

**Authors:** Mir Henglin, Maryam Ghareghani, William Harvey, David Porubsky, Sergey Koren, Evan E. Eichler, Peter Ebert, Tobias Marschall

## Abstract

Haplotype information is crucial for biomedical and population genetics research. However, current strategies to produce *de-novo* haplotype-resolved assemblies often require either difficult-to-acquire parental data or an intermediate haplotype-collapsed assembly. Here, we present Graphasing, a workflow which synthesizes the global phase signal of Strand-seq with assembly graph topology to produce chromosome-scale *de-novo* haplotypes for diploid genomes. Graphasing readily integrates with any assembly workflow that both outputs an assembly graph and has a haplotype assembly mode. Graphasing performs comparably to trio-phasing in contiguity, phasing accuracy, and assembly quality, outperforms Hi-C in phasing accuracy, and generates human assemblies with over 18 chromosome-spanning haplotypes.

## Background

Many eukaryotic organisms are diploid, and carry two sets of pairwise-similar chromosomes, with one set inherited from each parent. Consequently, separately assembling the two copies of each chromosome is necessary to fully characterize an individual’s genome. Each version of a chromosome inherited from a parent is called a *haplotype*. The process of assigning the two alleles of a heterozygous variant to their corresponding haplotype is termed *phasing*.

Haplotype-resolved genome assemblies provide crucial insights into studies of disease, evolution, and biodiversity by revealing segregation patterns of alleles within and between haplotypes [1]. Medically important genes and genomic regions, such as the major histocompatibility complex and *APOE* gene, exhibit compound heterozygosity, where alleles carried on the same haplotype produce a phenotype different than when those same alleles are carried on different haplotypes [2,3]. Haplotype-resolved assemblies support research on evolution, gene flow, demography, gene expression, and conservation biology [4–6], where knowledge of haplotype-specific combinations of genomic variants can be of crucial importance.

Despite their utility, it remains a major challenge to produce haplotype-resolved genome assemblies for diploid organisms. The ability of an assembler to phase genomic variation is directly tied to the length of the reads used to construct the assembly. As any single read originates from a single haplotype, any read that spans multiple heterozygous variants forms a “local” haplotype which can, in principle, be stitched into longer haplotype segments through the assembly of overlapping reads [2]. However, in practice this process is affected by both sequencing errors and ambiguities due to repetitive sequence. Consequently, advances in long-read genome sequencing technologies have led to improved genome assemblies, as reads lengths are now long enough to span a greater range of repetitive DNA variation [7]. Pacific Biosciences (PacBio) High-Fidelity (HiFi) reads [8] are 15–20 kb in length and have an error rate similar to accurate short-read sequencing, and Oxford Nanopore Technologies (ONT) Ultra-long reads [9] can achieve lengths > 100 kbp, which is long enough to span the majority of repeats found in human DNA. However, these read lengths are still too short to produce fully haplotype-resolved assemblies, even for assemblers utilizing combinations of long-read sequencing technologies [10,11]. Further computational steps and data sources beyond those employed in a “standard” genome assembly workflow are required in order to construct fully phased haplotypes [1,12–14].

When phasing with short, noisy, or low-coverage reads, reference-mapping-based methods are commonly used. Many phasing tools, such as WhatsHap [15], HapCol [16], HapCut2[17], MarginPhase [18], and LongPhase [19] utilize this strategy, where reads are first aligned to a reference genome and genomic variants are called. Subsequently, the variants are used to separate reads by haplotype for haplotype-specific assembly. The reference mapping approach is necessarily subjected to reference bias, and can therefore fail when variant calling is challenging due to unreliable alignment of reads to the reference, which occurs due to repetitive sequence or when the reference and sample differ in large structural variation [20]. Reference bias can be avoided by first constructing an unphased *de-novo* assembly to serve as the reference genome for genomic variant calling and phasing. This *de-novo* reference strategy is employed by the phasebook assembler [21], PGAS[22] and DipASM[22,23], where the latter two additionally leverage the long-range haplotype signal from Strand-seq [24,25] and Hi-C [26] data respectively to improve the haplotypes constructed with this strategy. However, the *de-novo* reference, being yet unphased, is a mosaic reference produced by collapsing sequence from both haplotypes together, which can introduce switch errors, false duplications, and nucleotide consensus errors [1,27–30].

When parental data is available, trio-binning can be used to assemble haplotypes without use of a reference genome. Trio-binning approaches use parental reads to identify “hap-mers”, k-mers unique to the maternal and paternal haplotypes, to label and partition reads before assembly [31]. Because trio-binning is reference-free, it avoids the errors introduced through the creation of a collapsed assembly. However, binning of reads before assembly is vulnerable to false duplications and fragmentation [32] and can be limited in its ability to phase repetitive or homozygous regions, which have few haplotype-specific k-mers [31,33].

Instead of binning reads by haplotype prior to assembly, performing the phasing directly on the assembly graph has emerged as an attractive strategy. Graph-based phasing typically combines the phase signal inherent to an assembly graph with an additional source of phase information [12,32,34], avoiding the errors introduced by the binning of reads before assembly while usually yielding larger phasing blocks. Typically, long-range phasing information from trio or Hi-C is aligned to the graph and synthesized with the graph topology to construct haplotypes. Long read assemblers such as hifiasm [10], Verkko [11], and Shasta [33], all natively support trio and Hi-C data integration, and independent modules which employ trio or Hi-C graph-based phasing, such as GreenHill [35] and GFAse [33], have recently emerged. These modules are designed to integrate with a wide range of assemblers, and can provide graph-based phasing capabilities to diverse workflows.

Trio-based assemblies have typically served as the gold standard for phased assembly, and trio assemblies from hifiasm and Verkko are currently the highest-quality assemblies that can be produced. However, the difficulty and expense of acquiring and sequencing three individuals’ genetic information limits trio-binning’s widespread application, and provokes interest in single-sample methods, such as as those leveraging Hi-C or Strand-seq, which can produce phased assemblies using only material from the sample of interest. Hi-C is a commercially available sequencing technology that captures chromatin conformation information [36]. Because a piece of DNA is far more likely to physically interact with itself than any other molecule, Hi-C’s ability to provide information on the physical proximity of DNA segments can be used to determine which variants originate from the same haplotype [23,37]. However, Hi-C only provides a local phase signal, the strength of which diminishes with distance, in contrast to the global phase signal provided by trio. Strand-seq is a short-read, single-cell sequencing method that generates sequencing libraries derived from only one DNA strand from each chromosome [24]. This is achieved by using a thymidine analog, bromodeoxyuridine (BrdU), to target and remove the nascent DNA strand during a round of cell division. Like trio, Strand-seq provides global phase signal, which, when combined with its status as single-sample technology, makes it an attractive target for method development.

### Contribution

We present Graphasing, a Strand-seq alignment-to-graph-based phasing and scaffolding workflow that assembles telomere-to-telomere (T2T) human haplotypes using data from a single sample. Graphasing leverages a robust cosine similarity clustering approach to synthesize global phase signal from Strand-seq alignments with assembly graph topology, producing accurate haplotype calls and end-to-end scaffolds. We built assemblies for the NA24385 (HG002) and HG00733 genomes using Graphasing with the Verkko and hifiasm assemblers and compared the quality of the haplotypes with those constructed by native trio and Hi-C mode, and show that our method produced the highest-quality single-sample assemblies, which match or exceed trio-phasing in contiguity, phasing accuracy, and assembly quality.

Graphasing is implemented using the open source workflow language, Snakemake [38]. The pipeline takes as input an assembly graph in .gfa format and a set of Strand-seq libraries in .fasta format, and outputs a haplotype partition of the assembly graph, which can readily be used by assembly tools to produce a final assembly, as well as Strand-seq annotations that can facilitate further downstream analysis. Graphasing is available at https://github.com/marschall-lab/strand-seq-graph-phasing

## Results

### Graph-phasing method

Graphasing phases an assembly graph produced by an assembly tool such as Verkko or hifiasm. The Graphasing workflow can be summarized in five main steps:

1. Alignment of Strand-seq reads to assembly unitigs (Figure 1a),
2. Clustering of unitigs by chromosome (Figure 1b),
3. Correction of misoriented unitigs (Figure 1c),
4. Pooling of haplotype informative reads to shade the assembly graph (Figure 1d),
5. Threading of haplotypes through the shaded graph to phase and scaffold the assembly (Figure 1d).

**Figure 1.**
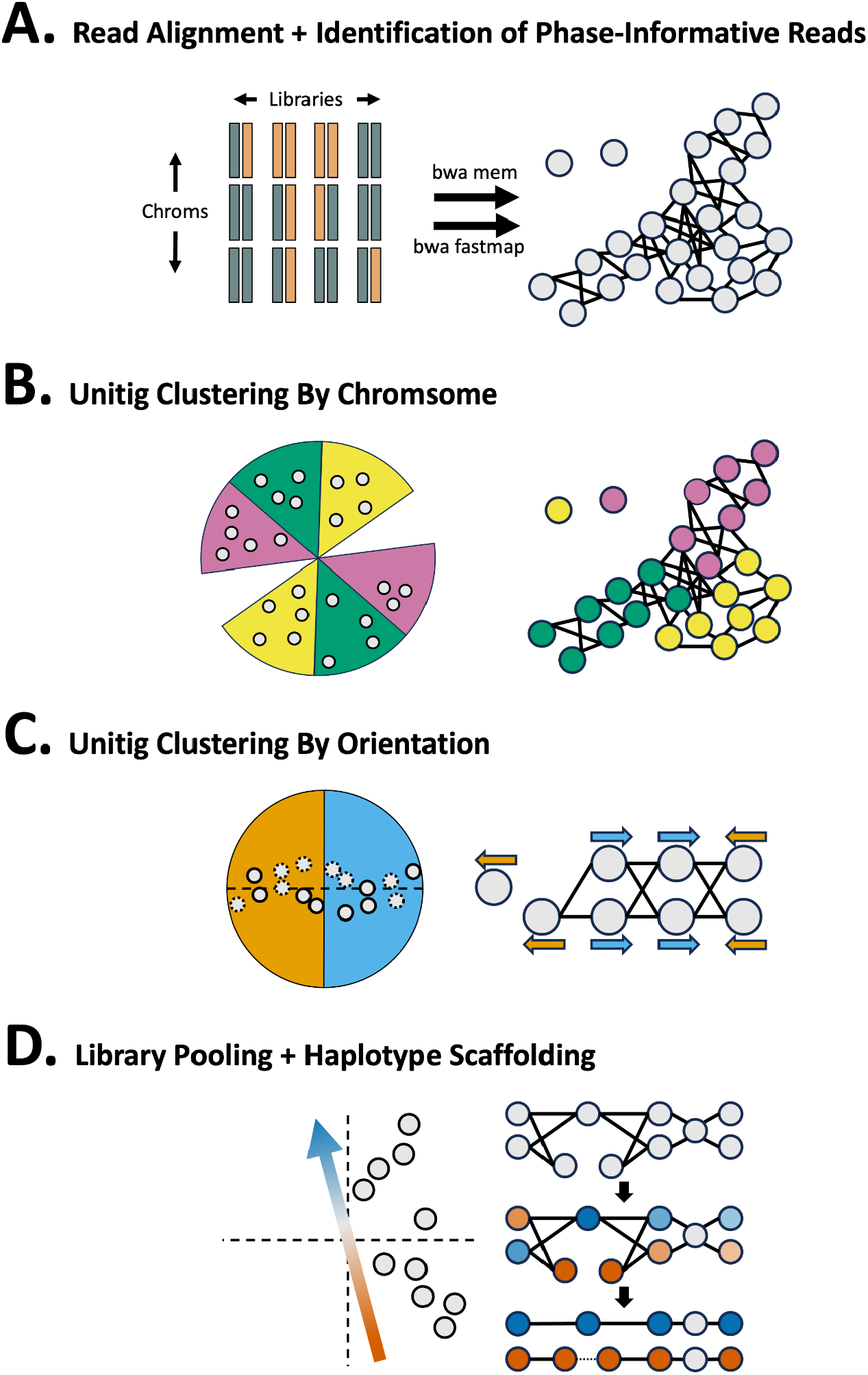
Pipeline overview. **A.** Reads from Strand-seq libraries are aligned to graph unitigs (gray circles) using ‘bwa mem’ and ‘bwa fastmap’.‘bwa fastmap’ alignments are used to identify haplotype informative reads, which are used for step “D” **B.** Unitigs (grey points) are clustered using a cosine-similarity based agglomerative clustering strategy. **C.** Unitigs (solid outline) and their flipped inverses (dotted outline) are used to correct misoriented unitigs. Unitigs in opposite orientation form a bisected structure that is captured with cosine-similarity clustering. **D.** The vector capturing the haplotype-informative libraries (left) is used to pool Strand-seq libraries and produce a haplotype shading of the assembly (right, middle). Rukki is run on the shaded graph to produce haplotype calls and scaffolds (right, bottom). Tangles and gaps are bridged, as indicated by the dotted line in the red haplotype.

Alignment of Strand-seq reads back to the genome convey global haplotype signal through the direction of the alignments [22,39–44] (Figure S1). However, though all reads can be used for clustering and misorientation correction, only reads aligning to unique sequence in the assembly carry phase signal, and these phase-informative reads are identified after alignment (Figure 1A). Unitig clustering by chromosome is performed using an agglomerative cosine-similarity clustering strategy (Figure 1B). Next, a hierarchical cosine-similarity clustering strategy is applied to identify misoriented unitigs in each chromosome cluster (Figure 1C). Finally, the phase-informative reads are pooled to produce a haplotype shading of the assembly graph (Figure 1D). Rukki [11] then threads the shaded graph to produce haplotype calls and scaffolds, which can bridge tangles and gaps in the assembly. Verkko directly accepts the output scaffolds as input to produce a phased assembly, while for hifiasm, phasing information is communicated by using the haplotype calls to construct k-mer databases that are passed to trio-mode assembly. Details of each step are described in the <u>Methods</u> section.

### Phasing Method Comparison

We compared the performance of Strand-seq based Graphasing to the results of the native trio and Hi-C phasing modes of Verkko and hifiasm. Assemblies were constructed for the NA24385 and HG00733 samples using the Verkko (v. 1.4.1) and hifiasm (v. 0.19.6) phasing pipelines. Hybrid assembly graphs were constructed with 118.1x coverage PacBio HiFi CCS reads [8] and 34.3x (6.3x >100kbp) coverage Oxford Nanopore Technologies (ONT) reads [9] for NA24385, and with 68.3x coverage PacBio HiFi CCS reads and 51.0x (32.8x >100kb) coverage Oxford nanopore for HG00733. Trio phased assemblies were constructed with parental short-read Illumina data at 30x coverage. Graphasing assemblies were constructed by inputting the unphased assembly graphs, using 192 libraries for NA24385 and 115 libraries for HG00733. Unitigs shorter than 50kbp were filtered out before phasing to reduce noise. Though these short unitigs made up a large fraction of the assembly by number, they represented at most 2.5% of the total sequence of a given unphased assembly (Table S1). It is important to note that Strand-seq and Hi-C phasing produce Haplotypes with Parentage Unknown (HaPUs), meaning that while each contig is haplotype-resolved, the parent-of-origin is unknown, unless further methods are employed [45]. Accordingly, evaluation was performed on both haplotypes together as a single assembly for each combination of assembler and sequencing technology. Here we present results for hybrid assemblies, but Graphasing can also produce high-quality haplotypes from hifiasm HiFi-only (44.5x) assemblies, which are competitive with trio phased-assemblies for NA24385 (Table S4).

#### Contiguity

Assembly contiguity was evaluated using N50 and auN. N50 is the most commonly reported metric of contiguity, and is defined as the length of the shortest contig for which longer and equal-length contigs cover more than 50% of the assembly [45], while the auN is a weighted sum of all Nx values for x between 0 and 100 [46]. The hifiasm assemblies were not scaffolded, while Verkko produces scaffolds, and therefore the Verkko assemblies were evaluated both on the scaffolds, and on the resulting scaftigs after breaking scaffolds at gaps.

We found that all phasing methods produced highly contiguous assemblies, with hifiasm auN ranging from 93.1 Mbp to 130.9 Mbp, Verkko auN ranging from 85.0 Mbp to 132.9 Mbp, and Verkko scaffold auN ranging from 135.4 Mbp to 146.4 Mbp (Table 1). To evaluate that improvement gained through each phasing method, we compared each assembly against its unphased counterpart, and found that each assembly was substantially more contiguous, having an N50 and auN at least 4-times larger, with larger gains observed for NA24385, which had a less contiguous input. Notably, the NA24385 scaffolds were more contiguous than the HG00733 scaffolds despite a less contiguous input graph, which investigations attributed to the presence of “hairpin-capped broken bubbles” in the center of the largest HG00733 chromosomes which fragmented some of the Rukki scaffolds (Figure S2).

**Table 1.** Assembly Contiguity Statistics. Phased Verkko assemblies list two numbers: the contig statistic first, and the scaffold statistic in parentheses second.

| Sample | Assembler | Phasing | N50 (Mbp) | auN (Mbp) |
| --- | --- | --- | --- | --- |
| HG00733 | hifiasm Hybrid | Trio | 104.0 | 105.6 |
| HG00733 | hifiasm Hybrid | Strand-seq | 110.0 | 113.7 |
| HG00733 | hifiasm Hybrid | Hi-C | 133.3 | 130.9 |
| HG00733 | hifiasm Hybrid | Unphased | 6.9 | 8.1 |
| HG00733 | Verkko Hybrid | Trio | 119.3 (137.8) | 124.3 (135.4) |
| HG00733 | Verkko Hybrid | Strand-seq | 134.8 (140.3) | 131.8 (140.2) |
| HG00733 | Verkko Hybrid | Hi-C | 134.9 (140.3) | 132.9 (140.0) |
| HG00733 | Verkko Hybrid | Unphased | 26.3 | 28.3 |
| NA24385 | hifiasm Hybrid | Trio | 95.8 | 99.0 |
| NA24385 | hifiasm Hybrid | Strand-seq | 95.3 | 93.1 |
| NA24385 | hifiasm Hybrid | Hi-C | 103.8 | 106.8 |
| NA24385 | hifiasm Hybrid | Unphased | 1.9 | 3.6 |
| NA24385 | Verkko Hybrid | Trio | 80.0 (143.7) | 85.0 (145.1) |
| NA24385 | Verkko Hybrid | Strand-seq | 89.1 (137.8) | 95.4 (146.4) |
| NA24385 | Verkko Hybrid | Hi-C | 87.1 (135.6) | 88.1 (142.3) |
| NA24385 | Verkko Hybrid | Unphased | 3.4 | 6.5 |

#### Nx Curves

For additional insight, we plotted each assembly’s Nx curve [48], which is created by plotting all Nx values, and additionally compared them against two high-quality reference assemblies; NA24385 was compared against the Q100 Project v1.0 NA24385 assembly [49,50] and HG00733 was compared against the T2T v2.0 CHM13 assembly [51].

Inspecting the Nx curves (Figure 2), we see that the Verkko Nx curves are mostly equidistant from the reference along the entire length of the curve, roughly indicating equivalent phasing performance at all lengths in the assembly. In contrast, hifiasm assemblies are much closer to the reference curve on the left side of the plots than on the right, indicating a relative dip in contiguity after the very largest contigs. For hifiasm, Hi-C phasing constructed the most contiguous assemblies, with its Nx curve the highest along nearly the entire domain for both samples, while Strand-seq and Trio performed comparably with one another. For Verkko, all assemblies were comparable with one another, with the exception of the Hi-C scaffolds, whose curves dips below the Strand-seq and Trio curves for the shorter contigs.

**Figure 2.**
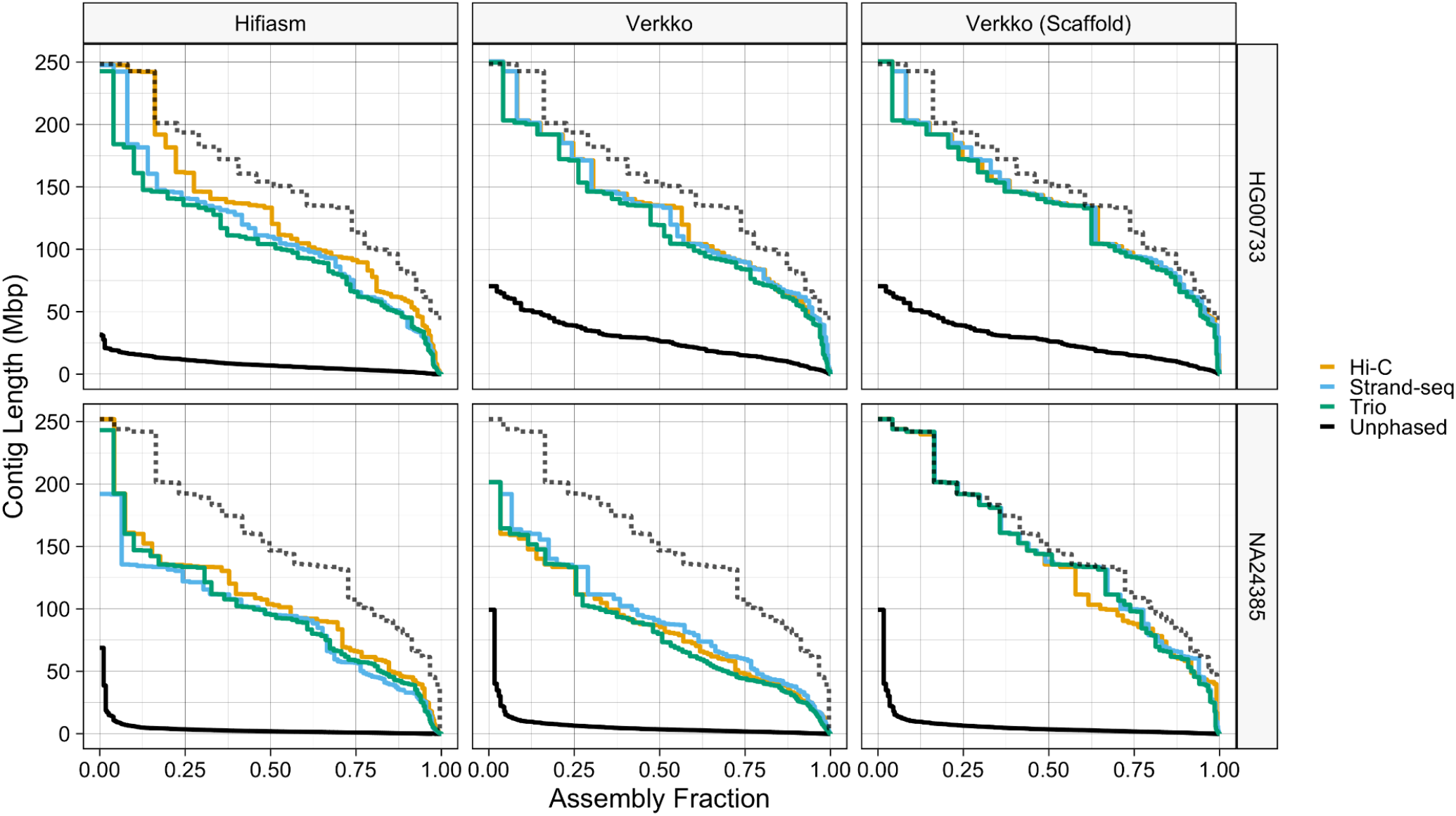
Nx curves. The dotted black line in each facet corresponds to the reference standards, which are the Q100 v1.0 assembly for NA24385, and the CHM13 v2.0 assembly for HG00733.

#### End-to-End Haplotypes

We further investigated each assembly for the number of end-to-end haplotypes. After using minimap2 [52,53] to align the assemblies to their respective references, the CHM13 v2.0 assembly for HG00733 and the Q100 v1.0 assembly for NA24385, three different properties were evaluated. If the summed alignment length was within 5% of the length of both the contig or scaffold and the reference chromosome, it was labeled “chromosome-scale”. If ‘seqtk telò [54] detected telomeric repeats at both ends of a contig or scaffold, it was labeled as having two telomeres. Finally, if a contig or scaffold mapped to the reference in one contiguous alignment, it was labeled “unbroken”. Unbroken alignments were only expected for the NA24385 assemblies, as they were aligned to a reference of the same genome. A contig or scaffold satisfying both of the first two properties was considered “chromosome-spanning”, while a contig or scaffold satisfying all three properties was considered to be “telomere-to-telomere” (T2T).

We found that for HG00733, all three Verkko assemblies performed comparably, while among the hifiasm assemblies, Hi-C had more chromosome-spanning contigs. (Table 2). For NA24385, Hi-C again performed the best among the hifiasm assemblies, while for Verkko, Graphasing produced a larger number of T2T and end-to-end contigs than trio phasing, despite having fewer chromosome-spanning scaffolds, while Hi-C produced the fewest chromsome-scale contigs and scaffolds. All Verkko NA24385 chromsome-spanning contigs were also T2T, while most hifiasm NA24385 chromsome-spanning contigs aligned to the reference in multiple pieces. Scaffolding greatly increased the number of chromosome-spanning sequences, and the Verkko assemblies each contained 15-25 chromosome-spanning scaffolds.

**Table 2.** End-to-end haplotype counts. Phased Verkko assemblies list two numbers: the contig statistic first, and the scaffold statistic in parentheses second.

| Sample | Assembler | Phasing | Chromosome-Scale (n) | Chromosome-Scale w/ Two Telomeres (n) | Chromosome-Scale w/ Two Telomeres & Unbroken (n) |
| --- | --- | --- | --- | --- | --- |
| HG00733 | hifiasm Hybrid | Trio | 15 | 8 | NA |
| HG00733 | hifiasm Hybrid | Strand-seq | 17 | 10 | NA |
| HG00733 | hifiasm Hybrid | Hi-C | 21 | 16 | NA |
| HG00733 | Verkko Hybrid | Trio | 17 (25) | 16 (22) | NA (NA) |
| HG00733 | Verkko Hybrid | Strand-seq | 21 (27) | 16 (20) | NA (NA) |
| HG00733 | Verkko Hybrid | Hi-C | 21 (26) | 17 (21) | NA (NA) |
| NA24385 | hifiasm Hybrid | Trio | 13 | 9 | 3 |
| NA24385 | hifiasm Hybrid | Strand-seq | 10 | 5 | 4 |
| NA24385 | hifiasm Hybrid | Hi-C | 16 | 11 | 7 |
| NA24385 | Verkko Hybrid | Trio | 4 (32) | 3 (25) | 3 (4) |
| NA24385 | Verkko Hybrid | Strand-seq | 10 (30) | 7 (18) | 7 (7) |
| NA24385 | Verkko Hybrid | Hi-C | 7 (27) | 5 (15) | 5 (5) |

#### Phasing Accuracy

Yak [32] was used to calculate the switch error rate and Hamming error rate. Yak utilizes parental sequence data to identify hap-mers and create a haplotype coloring of the assembly contigs and estimate switch and Hamming errors. To avoid inflation of the trio assemblies’ performance, the data used for error rate calculation was independent of the data used for trio phasing. For HG00733, hap-mers were identified from orthogonal parental Illumina sequencing data, and for NA24385, hap-mers were identified from the Q100 Project v1.0 assembly. The Q100 assembly is the highest quality NA24385 assembly publicly available, with an estimated error rate below 1 per 10 million bases [55]. For the Verkko assemblies, only scaffolds were evaluated in this and all subsequently described evaluations.

The assemblies produced by hifiasm and Verkko were generally high-quality and had a low error rate (Figure 3). For both samples, all phasing methods produced hifiasm assemblies with similar Hamming and switch error rates, with Hamming error rates around 0.9% and switch error rates around 0.85% for HG00733, and Hamming error rates around 0.20% and switch error rates around 0.13% for NA24385. Among the the Verkko assemblies, Graphasing and trio were the best performing for both samples, with switch and Hamming error rates below 0.85% for HG00733 and switch error rates below 0.09% and Hamming error rates below 0.07% for NA24385. The Verkko Hi-C assemblies, despite having a similar switch error rate as the other Verkko assemblies, had Hamming error rates about 1.5 and 5 times higher for HG00733 and NA24385 respectively, resulting from large, balanced switch errors (Figure 4). For NA24385, each Verkko haplotype had a switch error on average 0.06pp lower than the corresponding hifiasm haplotype which, though small in absolute terms, represents an approximate 2-fold difference in the switch error rate. A notable feature is that the NA24385 error rates are an order of magnitude less than the error rates of the HG00733 haplotypes. We believe that a large portion of the difference between the HG00733 and NA24385 error rates is due to the more accurate evaluation of the NA24385 haplotypes provided through the highly curated Q100 assembly, which suggests that the true error rates for the HG00733 assemblies may be lower than presented here.

**Figure 3.**
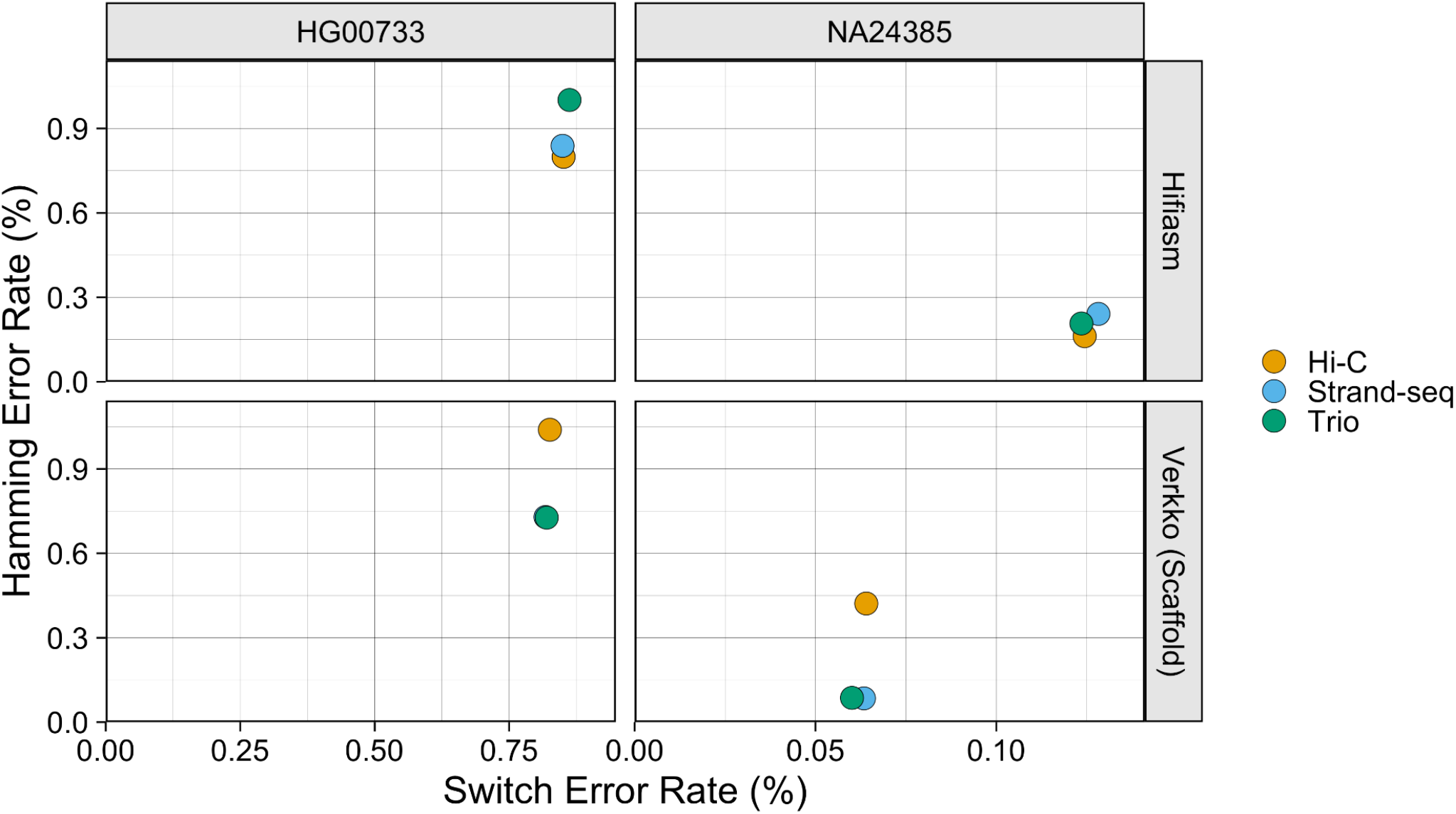
Haplotype Error Rate Scatter The X-coordinate of each point is the estimated switch error rate for a haplotype, and the y-coordinate is the estimated Hamming error rate. Points are colored by phasing data.

**Figure 4.**
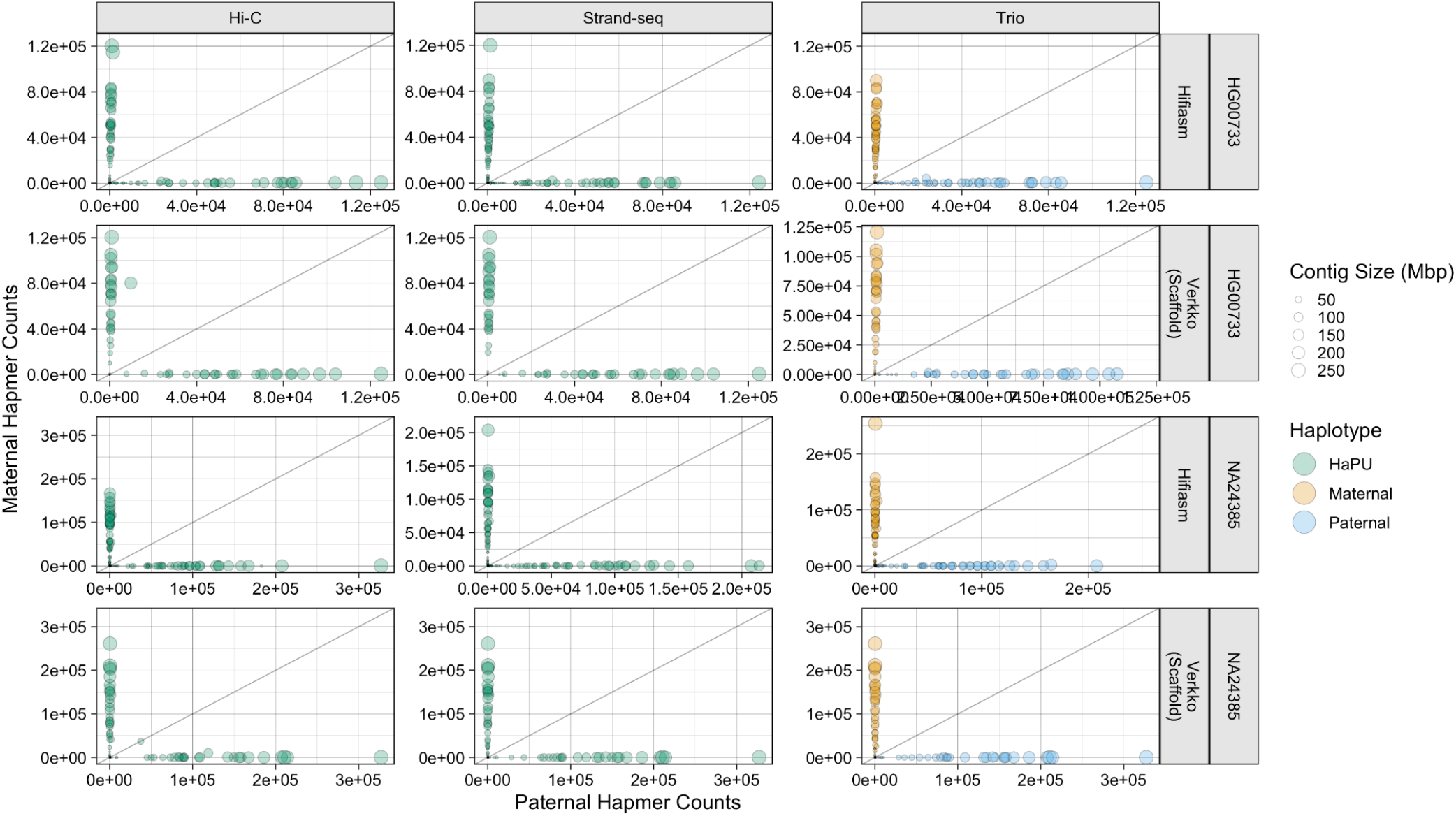
Hap-mer Blob Plots. For the NA24385 assemblies, only contigs aligning to autosomal chromosomes are plotted. The X- and Y-coordinate of each point is the number of hap-mers occuring on the contig, and the size of each point corresponds to contig length. Green points correspond to the Strand-seq and Hi-C HaPUs, while orange points correspond to the trio maternal haplotype, and blue points to the trio paternal haplotype. The grey line is the line of equality, where the number of hap-mers from either parent occurring on a contig is equal. The greater the phasing accuracy, the closer a blob is aligned to each axis.

To further investigate the phasing accuracy of the haplotypes, we produced hap-mer blob plots [56]. In a hap-mer blob plot, properly phased contigs, which contain hap-mers from only one parent, will be found on the X- or Y-axis. Any blob not on either axis contains a mixture of sequence from both parents, and contigs containing an equal mixture of parental hap-mers will be found on the gray line. Inspection of the blob plots revealed only the Verkko Hi-C assemblies had large, balanced switch errors, as the HG00733 and NA24385 assemblies each had single contig 185 and 42 Mbp in size respectively which strongly deviated from the axes (Figure 4). Smaller Hamming errors, represented through slight deviations from the axis can be observed in the other assemblies. Unitigs aligning to the X and Y chromosomes in the NA24385 assemblies received many more hap-mer alignments than unitigs aligning to the autosomes, and are plotted separately to preserve scale (Figure S3).

#### Consensus Quality

Consensus sequence quality value (QV) was estimated with Yak using orthogonal Illumina sequencing data for HG00733 and the Q100 v1.0 assembly for NA24385. Yak estimates the QV by comparing assembly k-mers to reference k-mers, with k-mers unique to the assembly presumed to be errors. Sequences shorter than 100kbp were filtered out before QV calculation.

All phasing methods produce high-quality assemblies with QV values >53 for all assemblies (Figure 5). For the Verkko assemblies, the Strand-seq and Hi-C haplotypes have similar QV scores, and both phasing methods slightly outperforming the trio assemblies. For the hifiasm assemblies, no haplotype strongly outperforms any other for HG00733, while Hi-C has a slight edge for NA24385.

**Figure 5.**
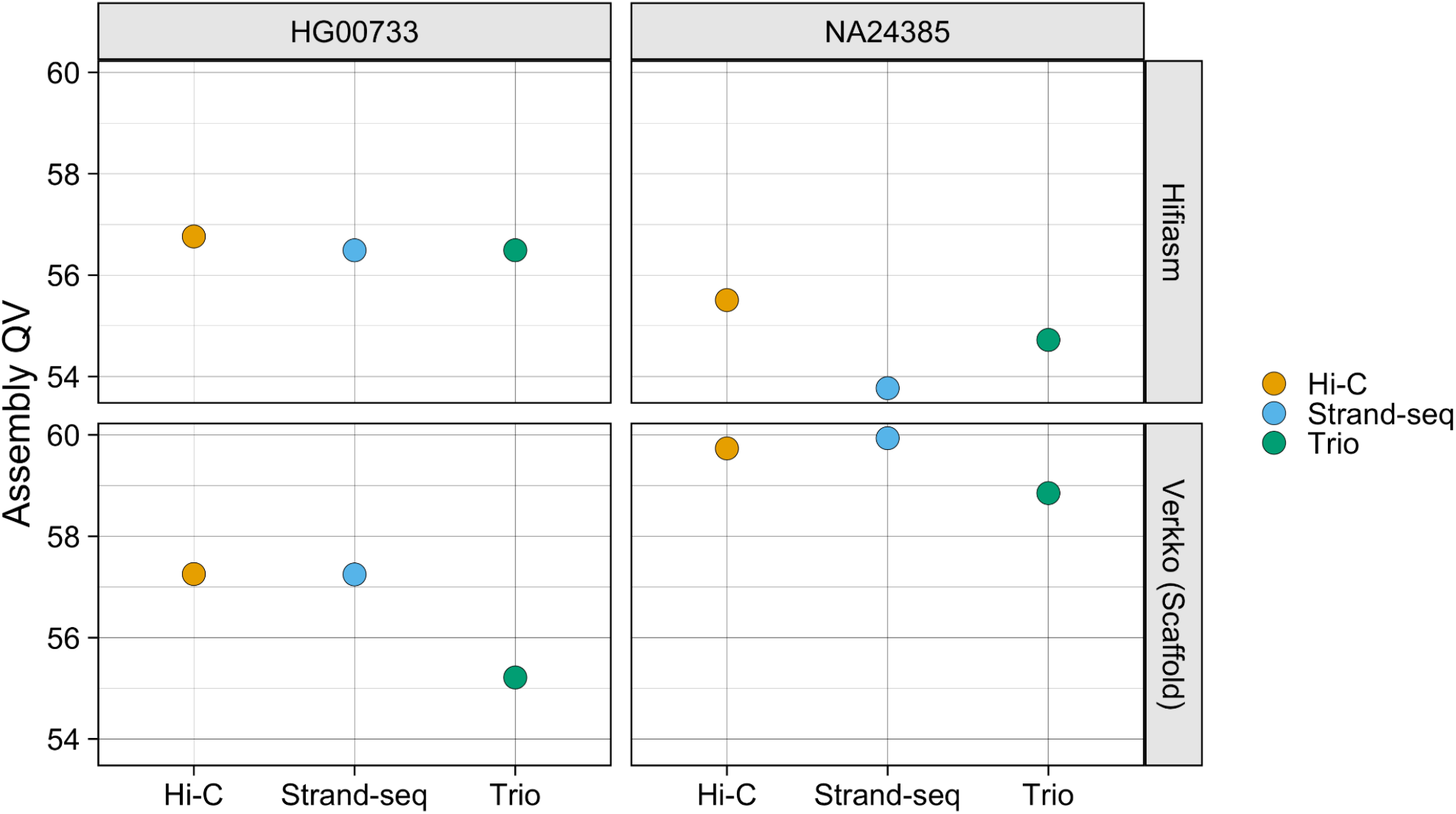
Assembly QV. Points are colored by phasing method.

#### Structural Misassemblies

Further evaluation was performed using paftools.js, a script included in the Minimap2 package [52]. ‘paftools misjoin’ counts gaps, inversions, and interchromosomal misjoins after aligning assembly contigs to a reference genome. The reference assemblies used were the T2T v2.0 CHM13 assembly [51] for HG00733, and the Q100 v1.0 assembly for NA24385, and alignment was performed with minimap2. ‘paftools.js misjoin’ was run with maximum gap size and minimum alignment block length thresholds of 1 Mbp. In our evaluation, we also examined the number of issues occurring entirely on unitigs aligning to acrocentric chromosomes, which are the most difficult to properly assemble and the most difficult to evaluate with alignment-based techniques.

Across all assemblies, the number of issues reported was low, with each assembly having no more than 17 detected events of a given category (Figure 6). Gaps were the most commonly reported event across all haplotypes, and mostly occurred on non-acrocentric chromosomes. More gaps were reported for the Verkko assemblies than for the hifiasm assemblies, as the scaffolded assemblies are naturally expected to contain gaps. Interchromosomal misjoins were the second most common event, and were reported only in unitigs aligning to acrocentric chromosomes. Due to the large amount of repetitive sequence within and between the acrocentric chromosomes, the interchromosomal misjoins may reflect a spurious call due to misalignment of the contigs to the reference [57]. Of the hifiasm assemblies, the Hi-C assemblies reported somewhat more misjoin events than assemblies phased by the other methods. Of the Verkko assemblies, results were nearly identical for all phasing methods for HG00733, while for NA24385, there is a clear ordering, with Hi-C performing the worst, and the trio assembly performing the best, reporting only two gaps and one misjoin. More issues were reported for the HG00733 assemblies than for the NA24385 assemblies, which may reflect genuine variation between the sample and CHM13 reference.

**Figure 6.**
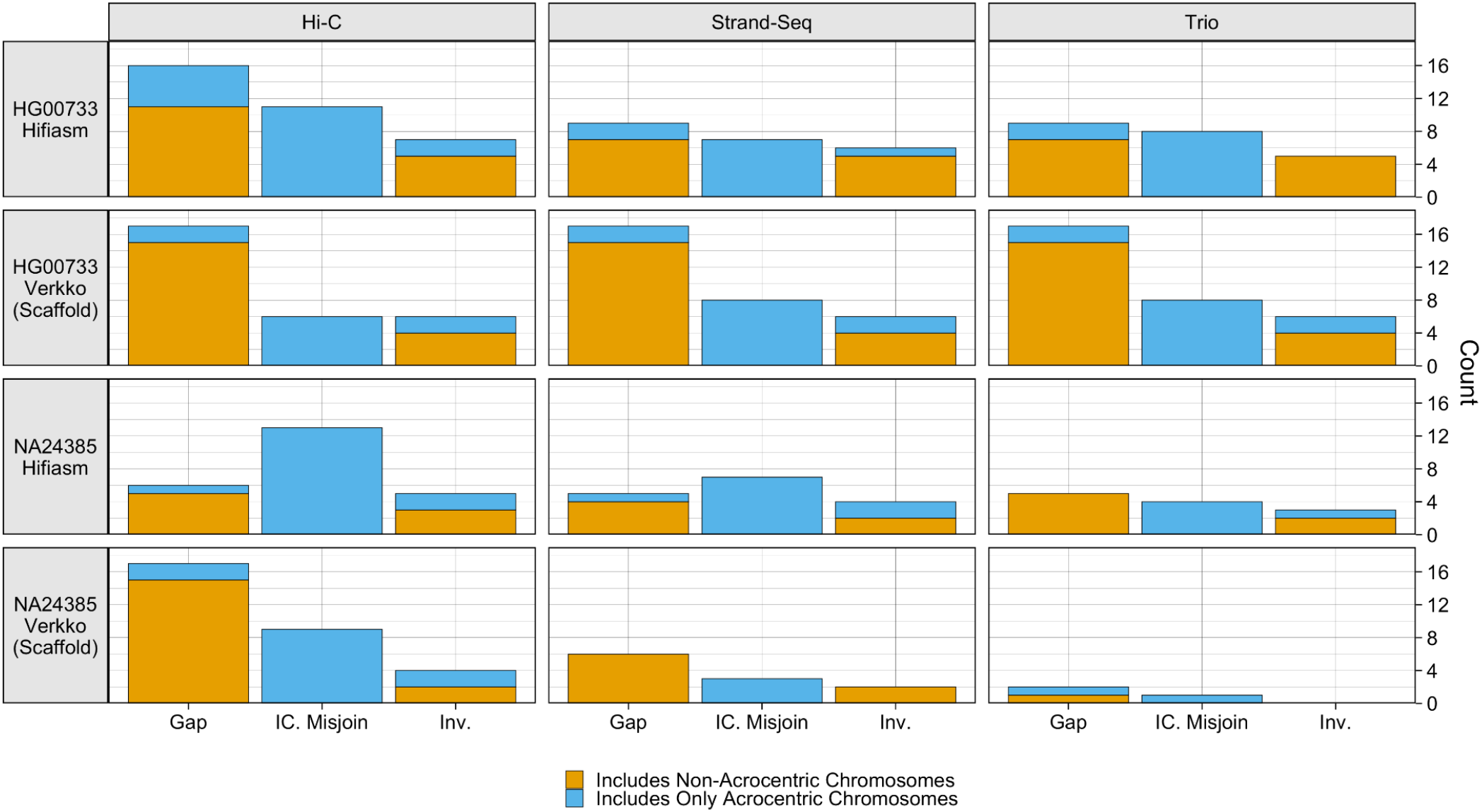
paftools.js misjoin Statistics: Three event categories are plotted: gaps, interchromosomal misjoins, and inversions. Each bar is colored blue according to the fraction of the misjoin type occurring entirely on acrocentric chromosomes (chromosomes 13, 14, 15, 21, 22).

#### Gene Completeness

‘paftools asmgene’ detects missing genes by aligning transcripts to both an assembly haplotype and a haploid reference and counting discrepancies in gene copy number. Subsequently, the percentage of genes that are multi-copy in the haploid reference but not in the assembly haplotype (%MMC) and the percentage of genes that are single-copy in the haploid reference but not in the assembly haplotype (%MSC) was computed. The reference assemblies used were the T2T v2.0 CHM13 assembly for HG00733, and the Q100 v1.0 assembly for NA24385, the transcripts came from Gencode v.44 protein-coding sequences [58], and alignment was performed with minimap2. For NA24385, each HaPU was compared against the reference haplotype according to the sex chromosome contained in the HaPU. Only full-length alignments with at least 99% identity were considered to label a gene as ‘present’ for the calculation of missing multi- and single-copy genes.

The assemblies showed low levels of gene missingness, with consistent patterns within samples (Figure 7). The NA24385 assemblies all had an MMC under 10% and MSC under 1.0%, and the HG00733 assemblies had an MMC under 12% and MSC under 2.5%. For HG00733, all phasing methods performed similarly for both assemblers, while for NA24385, Graphasing outperformed Hi-C, while trio-phasing, with an MMC below 1.51% and an MSC below 0.4% for both samples, showed by far the best results. However, the evaluation of the single-sample methods is deflated relative to trio, as Hi-C and Graphasing produce HaPUs but are evaluated against true haplotype references. This result also suggests that the gene completeness of the HG00733 assemblies, which were not evaluated against a reference of the same sample, is greater than the results presented.

**Figure 7.**
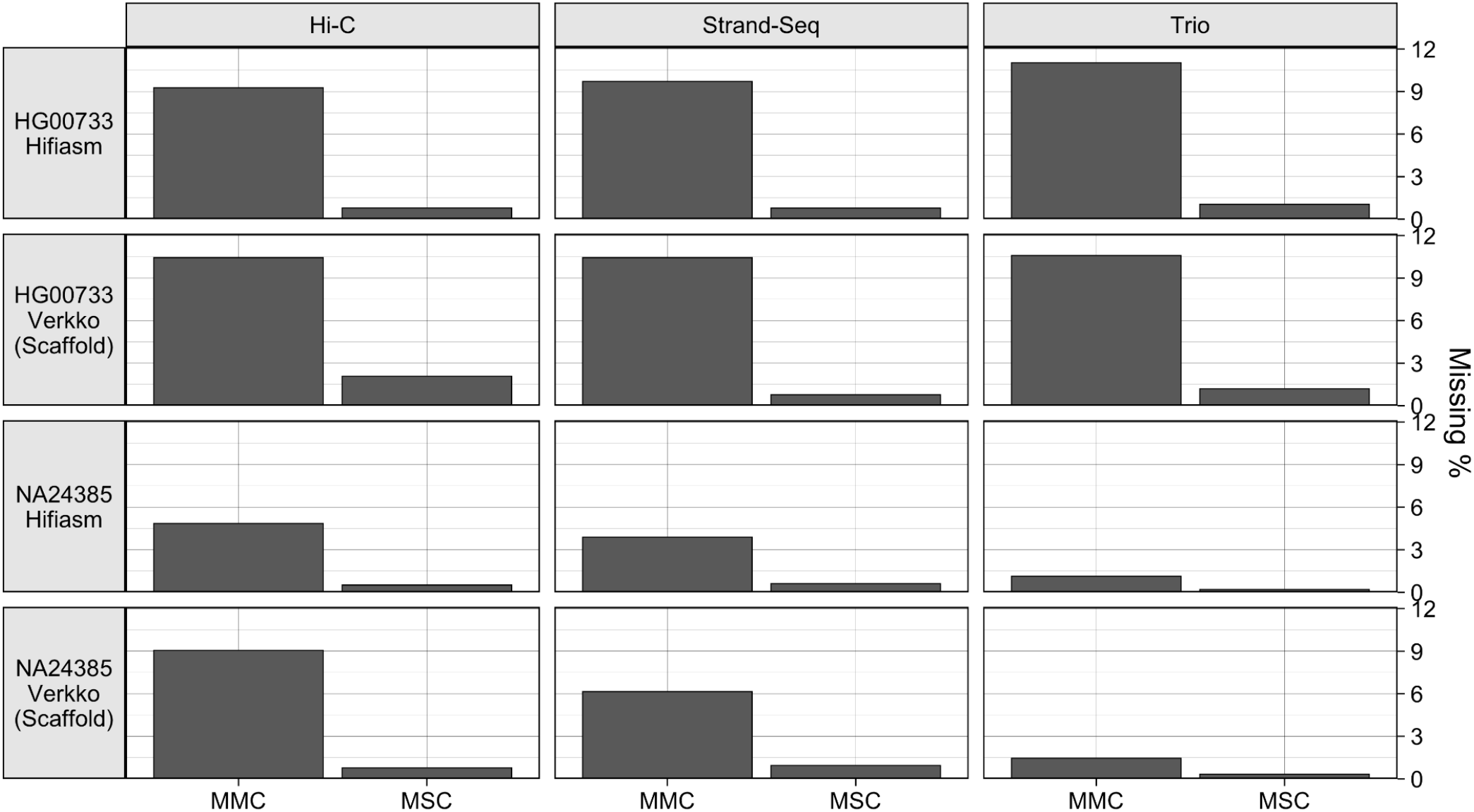
The fraction of missing multi-copy genes (MMC) and missing single-copy genes (MSC) calculated from paftools.js asmgene statistics.

### Strand-seq Library Titration

To evaluate the performance of Graphasing across varying Strand-seq input quality, a library titration experiment was run with the Verkko NA24385 sample. The 192 Strand-seq libraries had been previously annotated for quality, with 96 libraries labeled “high-quality” and libraries with a higher noise level and less clear phasing signal labeled “not-high-quality” (Table S2). With these annotations, 96 library sets were constructed by sampling without replacement 0%, 25%, 50%, 75%, or 100% of the libraries from the “high-quality” set, and sampling without replacement the remainder from the “not-high-quality” libraries. We sampled sets of size 96, as 96 is the number of libraries that is typically prepared in a single Strand-seq data preparation run. For the 0% and 100% library sets, as there is only one way to sample 0% or 100% of a set, there is only one sampled set. For each of the other percentages, four library sets were generated. Each sampled library set was then input to Graphasing, and the output haplotypes were compared against the HG002 v1.0 reference. Disagreement with the HG002 v1.0 reference was quantified as the percentage of the total assembly size, calculated using unitigs larger than the 50kbp input threshold, whose assignment does not match the reference, after aligning the unitigs to the reference. Results were quantified separately for the acrocentric chromosomes, which are the most difficult to assemble.

Our titration experiment showed that all library sets had high agreement with the HG002 v1.0 reference, with agreement values above 98% for the entire assembly, and above 93% for acrocentric chromosomes. (Figure 8). Disagreement averages slowly decrease with increasing library quality, to the minimum of 1.2% and 3.2% acrocentric disagreement for the 100% high-quality library set. Variance was higher for acrocentric-only disagreement, which was expected as acrocentrics contain large amounts of degenerate sequence that make alignments, and therefore Rukki haplotype assignments, unreliable. We additionally inspected the auN of the resulting scaffolds for each titrated set, and found that all samples achieved an auN within 16% of the reference auN, indicating that contiguity was also maintained across varying library compositions (Table S3). Our results indicate that high-quality phasing can be achieved across the entire range of Strand-seq input quality, as even a set of 96 low-quality libraries can still produce contiguous assembly with greater than 98% concordance with the HG002 v1.0 assembly for input unitigs longer than 50kbp.

**Figure 8.**
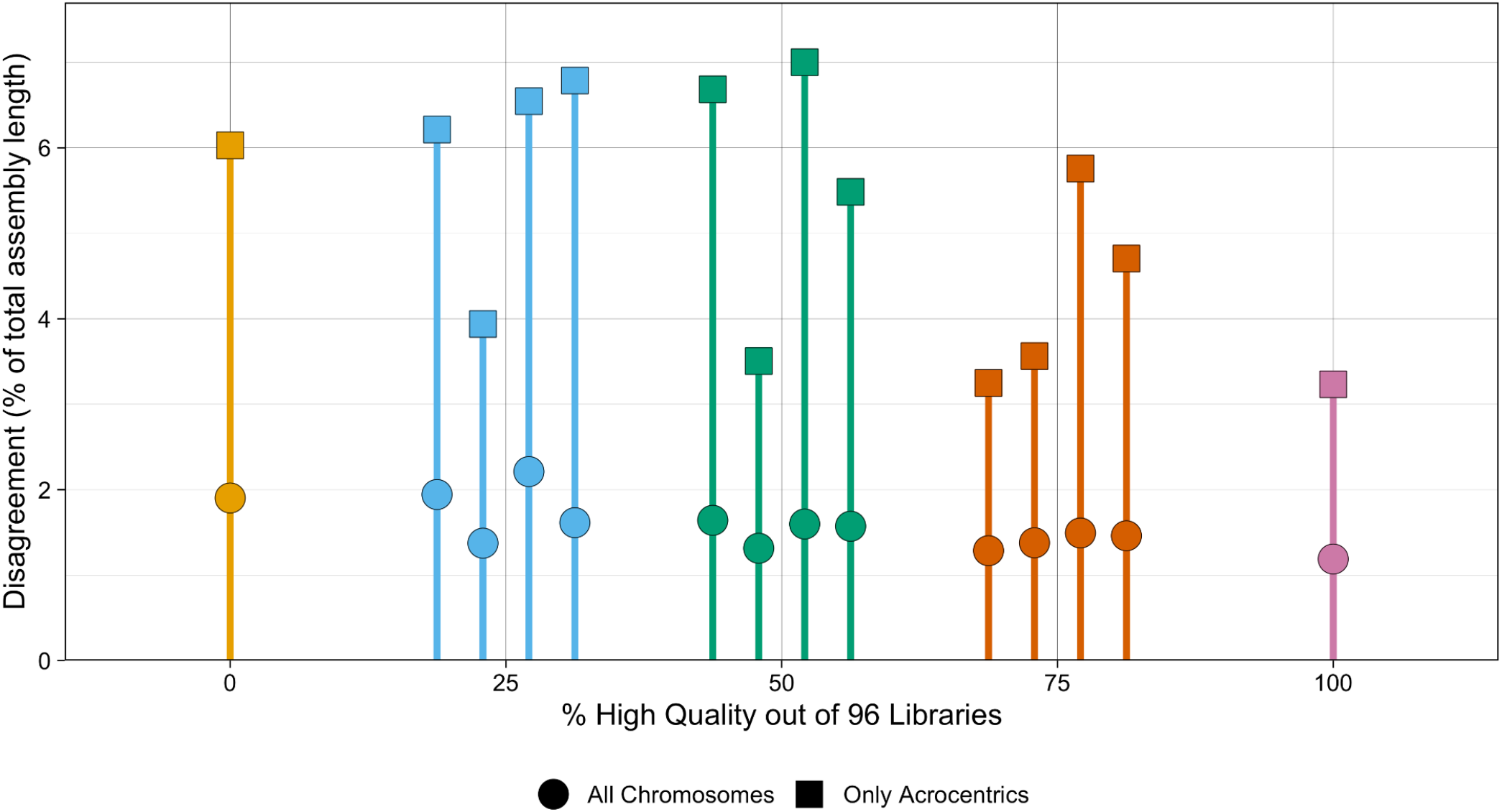
Disagreement between titrated and reference assemblies for NA24385. For each titrated Strand-seq library set, the haplotypes called by Rukki were compared to the reference haplotypes from the HG002 v1.0 assembly. Each color corresponds to a different fraction of high quality libraries sampled for the titrated library set, and shape corresponds to the inclusion or exclusion of unitigs aligning to the acrocentric chromosomes. Disagreement is quantified as the percent of the total length of the assembly for which haplotype calls disagree with the reference calls, calculated using unitigs longer than 50kbp.

### Runtime and memory usage evaluation

We evaluated the runtime and memory usage of Graphasing for all samples and assembly workflows (Table 3). Run time and peak memory usage of the tools were measured using the Snakemake “benchmark” decorator within Graphasing. Runtime and peak job memory usage were profiled on a computing cluster, with a standard cluster user profile. On a cluster, hifiasm runtime was around 8 and 11 hours and Verkko runtime was around 2.5 and 6 hours. The majority of the difference in runtime between assemblies was due to the contiguity of the input, with more fragmented assemblies taking longer to phase. Peak single job memory usage was at most 24GB for Verkko and at most 62GB for hifiasm across all runs. The greater peak job memory usage of the hifiasm assemblies came from creating k-mer databases with Yak. Regardless, the time and resources required are a small fraction of those used during a typical genome assembly workflow.

**Table 3.** Profiling Statistics.

| Sample | Assembler | Runtime (H:M) | Max Job Mem (GB) |
| --- | --- | --- | --- |
| HG00733 | Verkko Hybrid | 2:31 | 20 |
| NA24385 | Verkko Hybrid | 5:50 | 24 |
| HG00733 | hifiasm Hybrid | 8:02 | 62 |
| NA24385 | hifiasm Hybrid | 11:17 | 62 |

## Discussion

We introduced Graphasing, a workflow to phase genome assembly graphs, and compared its performance to the native Hi-C and trio phasing of Verkko and hifiasm for hybrid HiFI + ONT assemblies. Graphasing achieved performance comparable to that of trio phasing, as demonstrated through evaluation of contiguity, phasing accuracy, and assembly quality. In addition, we performed titration experiments to identify the range of input data quality under which Graphasing performs well. Input 96 library sets containing only high-noise libraries still achieved greater than 98% concordance with the HG002 v1.0 reference. Graphasing is modular and comprehensive, wrapping all operations from alignment to scaffolding, and adaptable to any assembler that outputs an assembly graph and has a phased assembly mode, making Graphasing widely applicable to different workflows.

In our evaluations, Verkko produced assemblies with similar contig-level contiguity as hifiasm for all assemblies. This result, when coupled with the fact that the Verkko scaffolds had similar or greater performance in assembly quality and phasing accuracy when compared to hifiasm contigs, represents an advantage for the Verkko assembler. Of the Verkko assemblies, all three phasing methods produced haplotypes with similar scaffold Nx curves and structural assembly quality. Among the single-sample methods, the Hi-C assemblies had higher phasing error, and a greater number of reported structural misassemblies and misassembled genes. Accordingly, we can state that the Verkko + Graphasing produced the highest-quality single-sample haplotypes.

The high contiguity of hybrid assemblies can present a unique methodological hurdle, despite the apparent decrease in phasing difficulty that comes from greater contiguity. Highly contiguous assemblies can contradict heuristics and challenge methods developed for more fragmented input. Another challenge of contiguous assemblies is when degenerate sequence is assembled alongside non-degenerate sequence onto a unitig; Degenerate genome regions receive alignments from multiple chromosomes, creating noise which can overwhelm phasing signal. The cosine-similarity based strategies utilized by Graphasing are robust to this noise and allow these challenging unitigs to be properly phased without preprocessing (Figure S4). Furthermore, Graphasing incorporates graph topology into the phasing process, allowing for a more robust phasing process that takes advantage of the highly contiguous graphs of hybrid assemblies. Nonetheless, some areas of the genome will remain challenging to phase due to the difficulty of accurately aligning short reads in those regions. Analysis of the phased haplotypes is also a challenge, as “ceiling effects” in quality analysis may pose an obstacle to accurately evaluating high-quality haplotypes.

Further downstream refinement and analyses of the phased assemblies, such as scaffolding acrocentric short arms or detection and analysis of inversions, can also be conducted with Strand-seq [59,60]. These analyses are facilitated by Strand-seq annotations computed by Graphasing. For example, one annotation identifies the phase-informative Strand-seq libraries, which allows for more informed investigation of apparent misjoins found in the assembly by allowing switch errors and misorientation events to be immediately distinguished from one another.

Graphasing is currently limited to diploid genomes. Extension to higher ploidy would require more input Strand-seq data as well as a significant rework of the core of the phasing workflow. Graphasing’s cosine-similarity approaches are effective for contiguous assemblies, but can struggle with more fragmented assemblies, as the approaches that work efficiently and effectively for contiguous assemblies can lead to trouble if there are many fragmented and degenerate unitigs in the input assembly. Strand-seq data can also be difficult to produce, given the need to isolate a single cell after a cycle of cell division. However, production of Strand-seq data is improving [61] and we consider the Graphasing pipeline to be an attractive assembly method especially for the production of reference-quality genomes when trio data is unavailable. Currently, Graphasing does not attempt to detect switch errors in the input assembly, and any switch errors present in the input assembly will propagate to the final haplotypes. Future iterations of the pipeline could include switch error detection and correction, a task for which Strand-seq already has proven successful [62].

## Conclusion

Graphasing is a Strand-seq-based phasing workflow that reconstructs chromosome-scale haplotypes from assembly graphs of diploid genomes. Comparison to gold-standard trio phasing shows that Graphasing achieves comparable performance across a range of evaluations of completeness, contiguity, and quality, and furthermore produces more complete and accurately phased assemblies than Hi-C phasing. Graphasing’s modular design allows it to be easily adapted to different assembly workflows. Both the phased genomes, as well as output Strand-seq annotations, facilitate further downstream analyses, such as missassembly detection, analysis of structural variants, and haplotype-specific gene analysis.

## Methods

### Aligning reads to assembly

While all reads can be used to cluster unitigs by chromosome, only a subset of reads convey haplotype information and are useful for phasing. Accordingly, reads are aligned to the assembly twice: once with bwa mem in paired-end mode [63], to derive the alignments used for clustering and orientation correction, and once with bwa fastmap [63] to identify the phasing-informative reads. bwa fastmap identifies super maximal exact matches (SMEMs), maximal exact matches that are not contained in any other maximal exact matches. Filtering to reads with only one SMEM filters out alignments to sequence that is present in multiple copies in the graph. This retains alignments to homozygous nodes and alignments that overlap heterozygous variation on diploid nodes. As bwa fastmap does not have a paired-end mode, reads are first merged with PEAR [64] to maximize utilized information. In cases where reads are not successfully merged, the first mate read is retained. Reads are homopolymer compressed before alignment for Verkko assemblies, as the Verkko assembly graph is also homopolymer compressed.

### Alignment Counting

Both the unitig clustering and phasing steps use only the aggregated counts of alignments in Watson and Crick orientation from each Strand-seq library. The processing steps before counting differs for each aligner. For the bwa mem alignments, duplicates are marked using sambamba [65] and then filtered out, along with supplementary, secondary, and improper alignments. bwa fastmap alignments are simply filtered to reads with only one SMEM. After filtering, the number of first-mate read alignments in Watson and Crick orientation from each Strand-seq library are counted for each unitig in the graph.

### Connected Components

The clustering step utilizes connected component information from the graph, following the heuristic that unitigs in the same connected component are more likely to have originated from the same chromosome than those in different connected components. However, unitigs from the five acrocentric chromosomes are expected to always be tangled together due to the high sequence similarity in the rDNA array. In an attempt to increase the utility of the connected component heuristic, Graphasing attempts to separate the acrocentric chromosomes before calculating the connected components. To do this, the largest connected component by number of base-pairs is first identified as the putative acrocentric cluster component. Subsequently, all nodes shorter than a threshold length, set by default to 50 kbp, are identified, and the largest tangle consisting solely of these short nodes on the putative acrocentric cluster component is labeled as the rDNA tangle. Nodes from the tangle, along with all edges connected to them, are then removed from the graph prior to calculation of connected components.

### Length Filtering

Unitigs shorter than an input threshold, which we set to 50kbp, are filtered out. The goal is to prevent short unitigs, which may either receive too few alignments to have a reliable signal or consist entirely of degenerate sequence, from adding noise that may disrupt accurate phasing of the assembly.

### Unitig Clustering

This step combines unitigs from homologous chromosomes into the same cluster. Unitig clustering can be broken into three stages: the first stage uses pre-processing functions from the contiBAIT R package [41] to filter out noisy libraries. Second, an initial clustering is created using a cosine-similarity based batched clustering strategy. Finally, this clustering is refined and completed using additional heuristics.

Strand-seq based chromosome clustering strategies [22,39,41,42] all rely on identifying shared patterns in the unitig strand state inherited across libraries. Each pair of homologous chromosomes inherits either an unmatched WC/CW strand state, or a matched WW/CC strand state for each Strand-seq library. Accordingly, all unitigs derived from the same pair of homologous chromosomes are expected to share strand states across Strand-seq libraries, making the unitig strand state a viable clustering signal. Though the exact strand state cannot be determined for each unitig and library, evidence for a matched or unmatched strand state can be quantified using the *strand state frequency* (SSF); let *w* and *c* be the number of Watson and Crick reads aligning to a unitig respectively. The SSF is defined as: (*w* − *c*)/(*w* + *c*). For a unitig with an equal number of Watson and Crick alignments, the SSF will be equal to 0, and when the alignments are all Watson or all Crick, the SSF will be 1 or −1 respectively. We therefore expect a matched strand state to produce SSF with a magnitude close to 1, and an unmatched strand state to result in an SSF close to 0. The SSF for a set of Stand-seq libraries is represented as a vector, where each component of the vector corresponds to a different Strand-seq library, and the value corresponds to the SSF for the library.

#### contiBAIT preprocessing

The contiBAIT preprocessing and clustering functions use a simple threshold to discretize the SSF and call strand states for each Strand-seq library. The preprocessing function then evaluates libraries for quality based on expected patterns in the strand states; because each unitig is expected to inherit matched and unmatched strand states in a 50/50 ratio across Strand-seq libraries, large deviations from this ratio indicate possible issues. Consequently libraries with too many unmatched strand states across unitigs, indicating possible failure of the Strand-seq chemistry, are discarded. Furthermore, unitigs and libraries with too few alignments to confidently call strand state are discarded.

#### Absolute Cosine Similarity Clustering

To understand why *absolute cosine similarity* is an appropriate metric for clustering unitigs by chromosome, we first consider the behavior of the SSF for ideal Strand-seq alignment data. Under ideal conditions, each unitig would have an SSF value of 0 for each library that inherited a unmatched strand state, and a value of 1 or −1 for each library that inherited a matched strand state. When considering the vector representation of the SSF, we see that *unitigs from the same chromosome will have vectors that all point along the same ideal vector, v*_*clust*_. *(Figure S4)*. The cosine similarity between two vectors *a* and *b* is defined as ||*a*||||*b*||*cos*θ where θ is the angle between *a* and *b*, and simplifies to *cos*θ if vectors *a* and *b* are unit-normalized. We thus see that the unit-normalized cosine similarity between two SSF vectors is maximized when they point in the same direction, making it an apt similarity metric for clustering. However, there is still a risk of misclustering misorientied unitigs, which can appear to originate from a different chromosome due to having a flipped value in matched strand state libraries. To account for possible misorientations, the absolute value of the cosine similarity is used, which clusters according to parallelism, regardless of direction.

Furthermore, cosine similarity is an appropriate metric for use with highly contiguous assemblies, where degenerate sequence becomes more likely to be assembled onto unitigs containing non-degenerate sequence. Repetitive genome regions receive alignments from multiple chromosomes, generating phasing noise. The cosine similarity based strategies utilized by Graphasing are robust to the noise generated by degenerate regions and allow challenging unitigs to be properly phased without preprocessing. This results from the noteworthy property of cosine similarity in that it reflects a relative, rather than absolute, comparison of the individual vector dimensions. Degenerate genome regions attract alignments from multiple chromosomes, and thus appear to have an unmatched strand state in every Strand-seq library. A degenerate region therefore shrinks each dimension of the SSF. However, because each non-zero component of the absolute SSF vector has a uniform magnitude, its normalized cosine similarity will not change if each dimension is shrunk by the same amount, making the metric robust to the effects of degenerate regions. An implicit assumption made by this metric is that inheriting an unmatched strand state in every library is impossible. While such an inheritance pattern is not ruled out by theory, it is extremely unlikely under the expectation that at least 96 Strand-seq libraries are input to the pipeline, 96 being the typical number of libraries generated in a single Strand-seq sequencing run. Therefore, we consider it safe to assume that an all unmatched strand-state inheritance pattern is the consequence of degenerate genomic regions.

#### Absolute Cosine Similarity Initial Clustering

A batched agglomerative clustering strategy is used to create the initial clustering. The unitigs are ranked based on coverage, and batched by quantile in groups of 1000. If fewer than 5 batches are created, then the unitigs are instead split into 5 quantiles. By clustering the batches in descending order of mean coverage, the clustering is seeded with “high-signal” unitigs, reducing variance. Additionally, batched clustering reduces clustering time by limiting the number of comparisons made at each clustering step.

The initial clustering algorithm consists of three main operations: cluster growing, cluster creation, and cluster merging. The first step is cluster growing, which begins by calculating the similarity between each unclustered unitig and each cluster, and if the largest similarity value exceeds a threshold value, then the unitig is added to the cluster. We define unitig-cluster similarity as the mean of the pairwise absolute cosine similarity calculated between the unclustered unitig and the unitigs in the cluster. Cluster growing is repeated until no similarities are greater than the specified threshold. At this point, the cluster creation step is triggered, which begins by calculating the similarity between all unclustered unitigs, and if the largest similarity value exceeds a threshold value, a new cluster is created from the two unitigs. If a new cluster is created, then the algorithm immediately loops back to cluster growing. Otherwise, the final operation, cluster merging, is triggered, where the similarity between clusters is calculated, and if the largest similarity value exceeds a threshold value, the clusters are merged. Here, we define the similarity between two clusters as the mean of the pairwise absolute cosine similarity calculated between the unitigs in each cluster. Cluster merging is first performed on each connected component, following the heuristic that unitigs on the same connected component have a higher chance of originating from the same chromosome, before a general merging step is performed. After cluster merging, the next batch of unitigs is added to the clustering process. At a given step, all unclustered unitigs from all batches added to the clustering process are considered for cluster growing or cluster creation. If all batches have been added, then the clustering routine ends.

#### Cluster Refinement

The first part of cluster refinement attempts to correct spurious clustering resulting from noisy unitigs. First, unitigs in clusters smaller than a specified minimum cluster size are relabeled as unclustered. Next, for each connected component, the fraction of the component basepairs labeled for each cluster is calculated, and unitigs belonging to clusters covering less than a specified threshold percentage, set by default to 2%, are relabeled as unclustered. These spurious clustering correction steps are necessary because minimal filtering of noisy unitigs is performed before clustering. Next, the number of clusters on each connected component is counted and, if there is only 1 cluster, the unclustered unitigs on the connected component are assigned to the cluster. Following this round of refinement, one more round of clustering, as described in “Absolute Cosine Similarity Initial Clustering”, is performed, using all unclustered unitigs. This is to attempt to assign the previously spuriously clustered unitigs to their correct clusters. Finally, a second round of refinement, which repeats the three steps described above, is performed.

#### Haploid Chromosome Clustering

To ensure that any small diploid regions attached to haploid chromosomes, such as the Pseudoautosomal region (PAR) are properly phased, unitigs from haploid chromosomes need to be identified, and their clusters merged before orientation correction and phasing. When the SSF is calculated using all reads, each unitig vector points along its corresponding *v*_*clust*_, corresponding to the chromosome from which it originated (Figure S4). When the SSF is calculated using only haplotype informative reads, each unitig vector will instead point along one of three different vectors corresponding to its haplotype membership: maternal, paternal, or homozygous. These three vectors lie in the “chromosome plane” spanned by *v*_*clust*_ and an orthogonal vector *v*_*phase*_, defined as the difference between the maternal and paternal haplotype vectors. (Figure S6). Haploid chromosome clusters do not contain unitigs from multiple haplotypes, however, and therefore all unitigs from a haploid chromosome cluster will instead continue to point along a single line regardless of which data is used for the SSF calculation. This gives a variance-based heuristic for identifying haploid chromosome clusters; after principal components analysis (PCA), haploid chromosomes are expected to have component 1 proportion of explained variance near 100%, and component 2 proportion of explained variance near 0%, in contrast to a more balanced ratio of values expected for diploid chromosome clusters. Accordingly, to identify haploid chromosome clusters, PCA is performed on each cluster after calculating the SSF vectors using only haplotype informative reads, and clusters with component 1 proportion of explained variance greater than 70% and component 2 proportion of explained variance less than 20% are labeled as homozygous chromosome clusters and merged.

### Cosine Similarity Unitig Orientation Correction

Before a shaded graph can be produced, misoriented unitigs need to be detected and corrected. In contrast to the chromosome clustering step, orientation correction uses non-absolute cosine similarity for clustering, as the SSF vectors of unitigs from the same chromosome but in opposite orientation will point in opposite directions along the same *v*_*clust*_, and therefore possess minimal cosine similarity. Accordingly, there is a natural bisection of the SSF vectors, with each cluster containing unitigs in the same orientation. To capture this structure, a two-cluster hierarchical clustering is performed. After clustering, the unitigs from an arbitrarily chosen cluster are corrected by “flipping” their orientation so that all unitigs in the cluster now have the same orientation. However, for graphs constructed with extremely high coverage data, the hierarchical clustering may capture structure other than unitig orientation. This risk arises from the fact that the unitigs from high-coverage hybrid assemblies can be extremely long and contiguous, such that a chromosome cluster may consist of only a few unitigs. In these cases, it is not unlikely that all unitigs may already be in the same orientation, meaning the bisected structure is not present for the hierarchical clustering to capture. To eliminate this risk, the clustering is performed on the unitigs together with a copy with the orientation “flipped”, which guarantees that unitigs in both orientations will be present when clustering. Afterwards, only the original version of each unitig is retained.

### Haplotype Informative Strand-seq Library Pooling

The sparse coverage of a typical Strand-seq library, generally ranging between 0.01x and 0.2x of the haploid genome [67], means that phase information from many libraries must be pooled to achieve a high quality result. Pooling haplotype informative reads requires two steps; identifying the unmatched strand state libraries, which are the libraries that convey phasing information, and properly assigning Watson and Crick labels to reads, such that all Watson reads are assigned to one haplotype and all Crick reads to the other haplotype (Figure S5). Previous work leverages identified SNVs [43] or homologous unitig pairs [68] to provide a supervising signal in a minimum error correction framework to achieve this goal. In contrast, Graphasing needs no external supervising signal to pool libraries, as all of the information needed for proper pooling can be derived from *v_phase_*. To calculate *v_phase_*, first, *v_clust_* is calculated as a size-weighted average of the orientation-corrected unitigs in a cluster. Next, PCA is performed on each cluster and the chromosome plane is inferred from the first two principal components. As described in “Haploid Chromosome Clustering”, it is expected that all unitig vectors will lie in the chromosome plane, and that *v_clust_* and *v_phase_* form an orthogonal basis for the plane. Because *v_clust_* and *v_phase_* are orthogonal, *v_phase_* can be inferred by projecting *v_clust_* into the chromosome plane, and then rotating the projected vector 90 degrees in the plane.

To understand how *v*_*phase*_ guides proper pooling, we make two observations. The first is that *v*_*phase*_ is expected to have a value of 1 or −1 if an unmatched strand state is inherited, and a value of 0 if a matched strand state is inherited, meaning that the non-zero components of *v*_*phase*_ identify the unmatched strand state libraries. The second is that WC and CW strand states will have opposite *v*_*phase*_ values of 1 and −1, meaning that two libraries with opposite sign assign assign reads of different orientation to a haplotype. Accordingly, proper pooling is achieved by swapping alignment counts in the libraries that have a *v*_*phase*_ value of −1, and then taking the dot product of each column of Watson or Crick alignment counts with *abs*(*v*_*phase*_). The pooled reads constitute a pair of haplotype marker counts for each unitig, whose values indicate the haplotype to which the unitig belongs. Unitigs specific to a given haplotype will have a high marker count for the corresponding haplotype, and a low marker count for the opposite haplotype, while homozygous unitigs are expected to be assigned a balanced number of markers for both haplotypes. Additionally, homozygous unitigs are expected to be assigned a much larger number of total markers than heterozygous unitigs, as all reads aligning to homozygous unitigs can be used for phasing.

#### Haploid Chromosome Phase Vector Correction

As mentioned above, *v_clust_* is calculated as a size-weighted average of the orientation-corrected unitig vectors within a cluster, which ensures that different levels of fragmentation between haplotypes, which could lead one haplotype to have more unitigs in the assembly than the other, do not skew the calculation. However, haploid chromosomes are often quite different in size, meaning that a size-weighted average will bias *v_clust_*, rotating it away from the ideal line of bisection between the two haploid chromosomes, biasing the inference of *v*_*phase*_. Accordingly, a balancing correction is performed for haploid chromosome clusters.

The correction is performed after calculation of *v_clust_* and *_*v*phase_* through the size-weighted average method described above. Each unitig vector is projected into the chromosome plane identified through the first two principal components, and then a change of basis is applied to express each projected vector in terms of *v_clust_* and *v_phase_*. After, the product of dimensions for each vector is calculated for each unitig, and the unitigs with the largest and smallest values are identified as “representatives.” The unitigs at these extremum are typically high-signal unitigs from either haploid chromosome. Afterwards, *v_clust_* and *v_phase_* are corrected by rotating both such that *v_clust_* bisects the two representative unitig vectors.

#### Haplotype Calling and Phased Consensus

Rather than call haplotypes for each unitig based on the pooled library counts alone, the counts are input to Rukki to be synthesized with graph topology and improve haplotype calls. The pooled counts create an initial shading of the graph, which is subsequently refined with Rukki graph-walking algorithms before a final haplotype call is output for each unitig. Rukki outputs haplotype scaffold paths in .gaf or .tsv format.

Currently, Graphasing generates files which may be input to the Verkko and hifiasm pipelines to generate a phased assembly .fasta. For Verkko, the haplotype scaffold paths .gaf can be directly input. For hifiasm, an indirect path must be taken to input the phasing information; a Yak kmer database is generated from each set of phased unitigs, which can be input to hifiasm trio mode to generate haplotype sequences.

## Declarations

### Availability of Data and Materials

Data analyzed in this study are all available from public repositories. NA24385 ONT data were acquired from the EPI2ME project and are available from the public Amazon S3 bucket s3://ont-open-data/. Data for which an ENA accession ID is listed can be accessed through the European Nucleotide Archive browser (https://www.ebi.ac.uk/ena/browser/home) while other data sources have a direct url (Table 4).

**Table 4.** Data availability.

| Sample | Data Type | ENA Accession/URL |
| --- | --- | --- |
| NA24385 | Strand-seq | url |
| HG00733 | Strand-seq | PRJEB12849 |
| NA24385 | Illumina (Phasing) | PRJNA477862 |
| HG00733 | Illumina (Phasing) | PRJNA477862 |
| HG00733 | Illumina (Evaluation) | PRJEB36890 (ERR3988823) |
| HG00732 | Illumina (Evaluation) | PRJEB31736 (ERR3241755) |
| HG00731 | Illumina (Evaluation) | PRJEB31736 (ERR3241754) |
| HG00733 | HiFi (Assembly) | url<br>url |
| NA24385 | HiFi (Assembly) | PRJNA731524<br>PRJNA813010 |
| HG00733 | ONT (Assembly) | url<br>url |
| NA24385 | ONT (Assembly) | url |
| What |  | URL |
| Gencode Transcripts |  | url |
| T2T-CHM13v2.0 Assembly |  | url |
| NA24385 Q100 v1.0 Assembly |  | url |
| hifiasm HiFi-only assembly files |  | NA24385_m64011_190830_220126.Q20.fastq.gz<br>NA24385_m64011_190901_095311.Q20.fastq.gz<br>NA24385_m64012_190920_173625.Q20.fastq.gz<br>NA24385_m64012_190921_234837.Q20.fastq.gz<br>NA24385_m54329U_201103_231616.Q20.fastq.gz |

## Acknowledgments

The authors thank Julian Lucas (UC Santa Cruz Genomics Institute) for his assistance in identifying data sources and providing guidance on Yak use and end-to-end haplotype identification.

## Authors’ Contributions

MG, DP, and TM designed an earlier version of a workflow to phase Hifi-based graphs using Strand-seq, which was implemented by MG. Based on this initial workflow, MH, PE, and TM designed the Graphasing method. MH implemented Graphasing. MH, WH and PE ran the assembly experiments. SK provided guidance on the integration with Verkko. MH prepared figures and wrote a draft manuscript. TM and PE edited the manuscript. MG and EEE helped with result interpretation and provided feedback on the manuscript. All authors read and approved the final manuscript

## References

1. Jarvis ED, Formenti G, Rhie A, Guarracino A, Yang C, Wood J, et al. Semi-automated assembly of high-quality diploid human reference genomes. Nature [Internet]. 2022;611:519–31. Available from: 10.1038/s41586-022-05325-5

2. Glusman G, Cox HC, Roach JC. Whole-genome haplotyping approaches and genomic medicine. Genome Med [Internet]. 2014;6:73. Available from: 10.1186/s13073-014-0073-7

3. Tewhey R, Bansal V, Torkamani A, Topol EJ, Schork NJ. The importance of phase information for human genomics. Nat Rev Genet [Internet]. 2011;12:215–23. Available from: 10.1038/nrg2950

4. Leitwein M, Duranton M, Rougemont Q, Gagnaire P-A, Bernatchez L. Using Haplotype Information for Conservation Genomics. Trends Ecol Evol [Internet]. 2020;35:245–58. Available from: 10.1016/j.tree.2019.10.012

5. Green RE, Krause J, Briggs AW, Maricic T, Stenzel U, Kircher M, et al. A draft sequence of the Neandertal genome. Science [Internet]. 2010;328:710–22. Available from: 10.1126/science.1188021

6. Cheng Y, Berg A, Wu S, Li Y, Wu R. Computing genetic imprinting expressed by haplotypes. Methods Mol Biol [Internet]. 2009;573:189–212. Available from: 10.1007/978-1-60761-247-6_11

7. Amarasinghe SL, Su S, Dong X, Zappia L, Ritchie ME, Gouil Q. Opportunities and challenges in long-read sequencing data analysis. Genome Biol [Internet]. 2020;21:30. Available from: 10.1186/s13059-020-1935-5

8. Wenger AM, Peluso P, Rowell WJ, Chang P-C, Hall RJ, Concepcion GT, et al. Accurate circular consensus long-read sequencing improves variant detection and assembly of a human genome. Nat Biotechnol [Internet]. 2019;37:1155–62. Available from: 10.1038/s41587-019-0217-9

9. Jain M, Koren S, Miga KH, Quick J, Rand AC, Sasani TA, et al. Nanopore sequencing and assembly of a human genome with ultra-long reads. Nat Biotechnol [Internet]. 2018;36:338–45. Available from: 10.1038/nbt.4060

10. Cheng H, Jarvis ED, Fedrigo O, Koepfli K-P, Urban L, Gemmell NJ, et al. Haplotype-resolved assembly of diploid genomes without parental data. Nat Biotechnol [Internet]. 2022;40:1332–5. Available from: 10.1038/s41587-022-01261-x

11. Rautiainen M, Nurk S, Walenz BP, Logsdon GA, Porubsky D, Rhie A, et al. Telomere-to-telomere assembly of diploid chromosomes with Verkko. Nat Biotechnol [Internet]. 2023; Available from: 10.1038/s41587-023-01662-6

12. Garg S. Computational methods for chromosome-scale haplotype reconstruction. Genome Biol [Internet]. 2021;22:101. Available from: 10.1186/s13059-021-02328-9

13. Sedlazeck FJ, Lee H, Darby CA, Schatz MC. Piercing the dark matter: bioinformatics of long-range sequencing and mapping. Nat Rev Genet [Internet]. 2018;19:329–46. Available from: 10.1038/s41576-018-0003-4

14. Nurk S, Koren S, Rhie A, Rautiainen M, Bzikadze AV, Mikheenko A, et al. The complete sequence of a human genome. Science [Internet]. 2022;376:44–53. Available from: 10.1126/science.abj6987

15. Patterson M, Marschall T, Pisanti N, van Iersel L, Stougie L, Klau GW, et al. WhatsHap: Weighted Haplotype Assembly for Future-Generation Sequencing Reads. J Comput Biol [Internet]. 2015;22:498–509. Available from: 10.1089/cmb.2014.0157

16. Pirola Y, Zaccaria S, Dondi R, Klau GW, Pisanti N, Bonizzoni P. HapCol: accurate and memory-efficient haplotype assembly from long reads. Bioinformatics [Internet]. 2016;32:1610–7. Available from: 10.1093/bioinformatics/btv495

17. Edge P, Bafna V, Bansal V. HapCUT2: robust and accurate haplotype assembly for diverse sequencing technologies. Genome Res [Internet]. 2017;27:801–12. Available from: 10.1101/gr.213462.116

18. Ebler J, Haukness M, Pesout T, Marschall T, Paten B. Haplotype-aware diplotyping from noisy long reads. Genome Biol [Internet]. 2019;20:116. Available from: 10.1186/s13059-019-1709-0

19. Lin J-H, Chen L-C, Yu S-C, Huang Y-T. LongPhase: an ultra-fast chromosome-scale phasing algorithm for small and large variants. Bioinformatics [Internet]. 2022;38:1816–22. Available from: 10.1093/bioinformatics/btac058

20. Masutani B, Suzuki Y, Suzuki Y, Morishita S. JTK: targeted diploid genome assembler. Bioinformatics [Internet]. 2023;39. Available from: 10.1093/bioinformatics/btad398

21. Luo X, Kang X, Schönhuth A. phasebook: haplotype-aware de novo assembly of diploid genomes from long reads. Genome Biol [Internet]. 2021;22:299. Available from: 10.1186/s13059-021-02512-x

22. Porubsky D, Ebert P, Audano PA, Vollger MR, Harvey WT, Marijon P, et al. Fully phased human genome assembly without parental data using single-cell strand sequencing and long reads. Nat Biotechnol [Internet]. 2021;39:302–8. Available from: 10.1038/s41587-020-0719-5

23. Garg S, Fungtammasan A, Carroll A, Chou M, Schmitt A, Zhou X, et al. Chromosome-scale, haplotype-resolved assembly of human genomes. Nat Biotechnol [Internet]. 2021;39:309–12. Available from: 10.1038/s41587-020-0711-0

24. Falconer E, Hills M, Naumann U, Poon SSS, Chavez EA, Sanders AD, et al. DNA template strand sequencing of single-cells maps genomic rearrangements at high resolution. Nat Methods [Internet]. 2012;9:1107–12. Available from: 10.1038/nmeth.2206

25. Sanders AD, Falconer E, Hills M, Spierings DCJ, Lansdorp PM. Single-cell template strand sequencing by Strand-seq enables the characterization of individual homologs. Nat Protoc [Internet]. 2017;12:1151–76. Available from: 10.1038/nprot.2017.029

26. Lieberman-Aiden E, van Berkum NL, Williams L, Imakaev M, Ragoczy T, Telling A, et al. Comprehensive mapping of long-range interactions reveals folding principles of the human genome. Science [Internet]. 2009;326:289–93. Available from: 10.1126/science.1181369

27. Church DM, Schneider VA, Graves T, Auger K, Cunningham F, Bouk N, et al. Modernizing reference genome assemblies. PLoS Biol [Internet]. 2011;9:e1001091. Available from: 10.1371/journal.pbio.1001091

28. Church DM, Schneider VA, Steinberg KM, Schatz MC, Quinlan AR, Chin C-S, et al. Extending reference assembly models. Genome Biol [Internet]. 2015;16:13. Available from: 10.1186/s13059-015-0587-3

29. Rhie A, McCarthy SA, Fedrigo O, Damas J, Formenti G, Koren S, et al. Towards complete and error-free genome assemblies of all vertebrate species. Nature [Internet]. 2021;592:737–46. Available from: 10.1038/s41586-021-03451-0

30. Kim J, Lee C, Ko BJ, Yoo DA, Won S, Phillippy AM, et al. False gene and chromosome losses in genome assemblies caused by GC content variation and repeats. Genome Biol [Internet]. 2022;23:204. Available from: 10.1186/s13059-022-02765-0

31. Koren S, Rhie A, Walenz BP, Dilthey AT, Bickhart DM, Kingan SB, et al. De novo assembly of haplotype-resolved genomes with trio binning. Nat Biotechnol [Internet]. 2018; Available from: 10.1038/nbt.4277

32. Cheng H, Concepcion GT, Feng X, Zhang H, Li H. Haplotype-resolved de novo assembly using phased assembly graphs with hifiasm. Nat Methods [Internet]. 2021;18:170–5. Available from: 10.1038/s41592-020-01056-5

33. Lorig-Roach R, Meredith M, Monlong J, Jain M, Olsen H, McNulty B, et al. Phased nanopore assembly with Shasta and modular graph phasing with GFAse. bioRxiv [Internet]. 2023; Available from: 10.1101/2023.02.21.529152

34. Garg S, Rautiainen M, Novak AM, Garrison E, Durbin R, Marschall T. A graph-based approach to diploid genome assembly. Bioinformatics [Internet]. 2018;34:i105–14. Available from: 10.1093/bioinformatics/bty279

35. Ouchi S, Kajitani R, Itoh T. GreenHill: a de novo chromosome-level scaffolding and phasing tool using Hi-C. Genome Biol [Internet]. 2023;24:162. Available from: 10.1186/s13059-023-03006-8

36. Beitel CW, Froenicke L, Lang JM, Korf IF, Michelmore RW, Eisen JA, et al. Strain- and plasmid-level deconvolution of a synthetic metagenome by sequencing proximity ligation products. PeerJ [Internet]. 2014;2:e415. Available from: 10.7717/peerj.415

37. Kronenberg ZN, Rhie A, Koren S, Concepcion GT, Peluso P, Munson KM, et al. Extended haplotype-phasing of long-read de novo genome assemblies using Hi-C. Nat Commun [Internet]. 2021;12:1935. Available from: 10.1038/s41467-020-20536-y

38. Köster J, Rahmann S. Snakemake--a scalable bioinformatics workflow engine. Bioinformatics [Internet]. 2012;28:2520–2. Available from: 10.1093/bioinformatics/bts480

39. Ghareghani M, Porubskỳ D, Sanders AD, Meiers S, Eichler EE, Korbel JO, et al. Strand-seq enables reliable separation of long reads by chromosome via expectation maximization. Bioinformatics [Internet]. 2018;34:i115–23. Available from: 10.1093/bioinformatics/bty290

40. Hills M, Falconer E, O’Neill K, Sanders AD, Howe K, Guryev V, et al. Construction of Whole Genomes from Scaffolds Using Single Cell Strand-Seq Data. Int J Mol Sci [Internet]. 2021;22. Available from: 10.3390/ijms22073617

41. O’Neill K, Hills M, Gottlieb M, Borkowski M, Karsan A, Lansdorp PM. Assembling draft genomes using contiBAIT. Bioinformatics [Internet]. 2017;33:2737–9. Available from: 10.1093/bioinformatics/btx281

42. Hills M, O’Neill K, Falconer E, Brinkman R, Lansdorp PM. BAIT: Organizing genomes and mapping rearrangements in single cells. Genome Med [Internet]. 2013;5:82. Available from: 10.1186/gm486

43. Porubsky D, Garg S, Sanders AD, Korbel JO, Guryev V, Lansdorp PM, et al. Dense and accurate whole-chromosome haplotyping of individual genomes. Nat Commun [Internet]. 2017;8:1293. Available from: 10.1038/s41467-017-01389-4

44. Porubský D, Sanders AD, van Wietmarschen N, Falconer E, Hills M, Spierings DCJ, et al. Direct chromosome-length haplotyping by single-cell sequencing. Genome Res [Internet]. 2016;26:1565–74. Available from: 10.1101/gr.209841.116

45. Lander ES, Linton LM, Birren B, Nusbaum C, Zody MC, Baldwin J, et al. Initial sequencing and analysis of the human genome. Nature [Internet]. 2001;409:860–921. Available from: 10.1038/35057062

46. Salzberg SL, Phillippy AM, Zimin A, Puiu D, Magoc T, Koren S, et al. GAGE: A critical evaluation of genome assemblies and assembly algorithms. Genome Res [Internet]. 2012;22:557–67. Available from: 10.1101/gr.131383.111

47. Akbari V, Hanlon VCT, O’Neill K, Lefebvre L, Schrader KA, Lansdorp PM, et al. Parent-of-origin detection and chromosome-scale haplotyping using long-read DNA methylation sequencing and Strand-seq. Cell Genom [Internet]. 2023;3:100233. Available from: 10.1016/j.xgen.2022.100233

48. Gurevich A, Saveliev V, Vyahhi N, Tesler G. QUAST: quality assessment tool for genome assemblies. Bioinformatics [Internet]. 2013;29:1072–5. Available from: 10.1093/bioinformatics/btt086

49. Zook JM, Hansen NF, Olson ND, Chapman L, Mullikin JC, Xiao C, et al. A robust benchmark for detection of germline large deletions and insertions. Nat Biotechnol [Internet]. 2020;38:1347–55. Available from: 10.1038/s41587-020-0538-8

50. Wang T, Antonacci-Fulton L, Howe K, Lawson HA, Lucas JK, Phillippy AM, et al. The Human Pangenome Project: a global resource to map genomic diversity. Nature [Internet]. 2022;604:437–46. Available from: 10.1038/s41586-022-04601-8

51. Rhie A, Nurk S, Cechova M, Hoyt SJ, Taylor DJ, Altemose N, et al. The complete sequence of a human Y chromosome. Nature [Internet]. 2023;621:344–54. Available from: 10.1038/s41586-023-06457-y

52. Li H. Minimap2: pairwise alignment for nucleotide sequences. Bioinformatics [Internet]. 2018;34:3094–100. Available from: 10.1093/bioinformatics/bty191

53. Li H. New strategies to improve minimap2 alignment accuracy. Bioinformatics [Internet]. 2021;37:4572–4. Available from: 10.1093/bioinformatics/btab705

54. Li H. seqtk: Toolkit for processing sequences in FASTA/Q formats [Internet]. Github; [cited 2024 Jan 26]. Available from: https://github.com/lh3/seqtk

55. HG002: A complete diploid human genome [Internet]. Github; [cited 2024 Jan 11]. Available from: https://github.com/marbl/HG002

56. Rhie A, Walenz BP, Koren S, Phillippy AM. Merqury: reference-free quality, completeness, and phasing assessment for genome assemblies. Genome Biol [Internet]. 2020 [cited 2024 Jan 17];21:1–27. Available from: https://genomebiology.biomedcentral.com/articles/10.1186/s13059-020-02134-9

57. Guarracino A, Buonaiuto S, de Lima LG, Potapova T, Rhie A, Koren S, et al. Recombination between heterologous human acrocentric chromosomes. Nature [Internet]. 2023;617:335–43. Available from: 10.1038/s41586-023-05976-y

58. Frankish A, Diekhans M, Ferreira A-M, Johnson R, Jungreis I, Loveland J, et al. GENCODE reference annotation for the human and mouse genomes. Nucleic Acids Res [Internet]. 2019;47:D766–73. Available from: 10.1093/nar/gky955

59. Porubsky D, Höps W, Ashraf H, Hsieh P, Rodriguez-Martin B, Yilmaz F, et al. Recurrent inversion polymorphisms in humans associate with genetic instability and genomic disorders. Cell [Internet]. 2022;185:1986–2005.e26. Available from: 10.1016/j.cell.2022.04.017

60. Sanders AD, Hills M, Porubský D, Guryev V, Falconer E, Lansdorp PM. Characterizing polymorphic inversions in human genomes by single-cell sequencing. Genome Res [Internet]. 2016;26:1575–87. Available from: 10.1101/gr.201160.115

61. Hanlon VCT, Chan DD, Hamadeh Z, Wang Y, Mattsson C-A, Spierings DCJ, et al. Construction of Strand-seq libraries in open nanoliter arrays. Cell Rep Methods [Internet]. 2022;2:100150. Available from: 10.1016/j.crmeth.2021.100150

62. Porubsky D, Sanders AD, Taudt A, Colomé-Tatché M, Lansdorp PM, Guryev V. breakpointR: an R/Bioconductor package to localize strand state changes in Strand-seq data. Bioinformatics [Internet]. 2020;36:1260–1. Available from: 10.1093/bioinformatics/btz681

63. Li H. Aligning sequence reads, clone sequences and assembly contigs with BWA-MEM [Internet]. arXiv [q-bio.GN]. 2013. Available from: http://arxiv.org/abs/1303.3997

64. Zhang J, Kobert K, Flouri T, Stamatakis A. PEAR: a fast and accurate Illumina Paired-End reAd mergeR. Bioinformatics [Internet]. 2014;30:614–20. Available from: 10.1093/bioinformatics/btt593

65. Tarasov A, Vilella AJ, Cuppen E, Nijman IJ, Prins P. Sambamba: fast processing of NGS alignment formats. Bioinformatics [Internet]. 2015;31:2032–4. Available from: 10.1093/bioinformatics/btv098

66. O’Neill K. Automated analysis of single cell leukemia data [Internet]. University of British Columbia; 2014 [cited 2023 Oct 3]. Available from: https://open.library.ubc.ca/soa/cIRcle/collections/ubctheses/24/items/1.0135595

67. Hanlon V, Porubsky D, Lansdorp P. Chromosome-length haplotypes with StrandPhaseR and Strand-seq [Internet]. The University of British Columbia; 2022. Available from: https://doi.library.ubc.ca/10.14288/1.0406302

68. Ghareghani M. Single-cell strand sequencing for structural variant analysis and genome assembly [Internet]. Universität des Saarlandes; 2022. Available from: https://publikationen.sulb.uni-saarland.de/handle/20.500.11880/34644

69. Chin C-S, Peluso P, Sedlazeck FJ, Nattestad M, Concepcion GT, Clum A, et al. Phased diploid genome assembly with single-molecule real-time sequencing. Nat Methods [Internet]. 2016;13:1050–4. Available from: 10.1038/nmeth.4035

