## supplement for "Phasing Diploid Genome Assembly Graphs with Single-Cell Strand Sequencing"

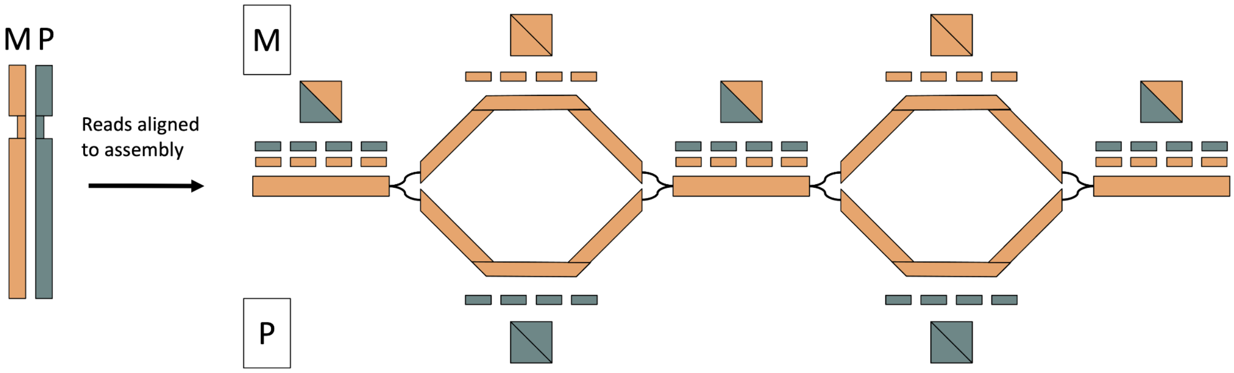


*Figure S1. Strand-seq Phasing Signal. A reads from a WC Strand-seq library (left) are aligned to the unitigs of a chromosome (right). Orange shapes correspond to Watson orientation, and green shapes correspond to Crick orientation. The graph unitigs and maternal reads are all in Watson orientation, while the paternal reads are all in Crick orientation. Consequently, sequence unique to the maternal haplotype will receive only "forward" alignments (orange box), while sequence unique to the paternal haplotype will receive only "reverse" alignments (green box). Sequence shared between the haplotypes receives a 50/50 mix of forward and reverse alignments (green\orange box)*

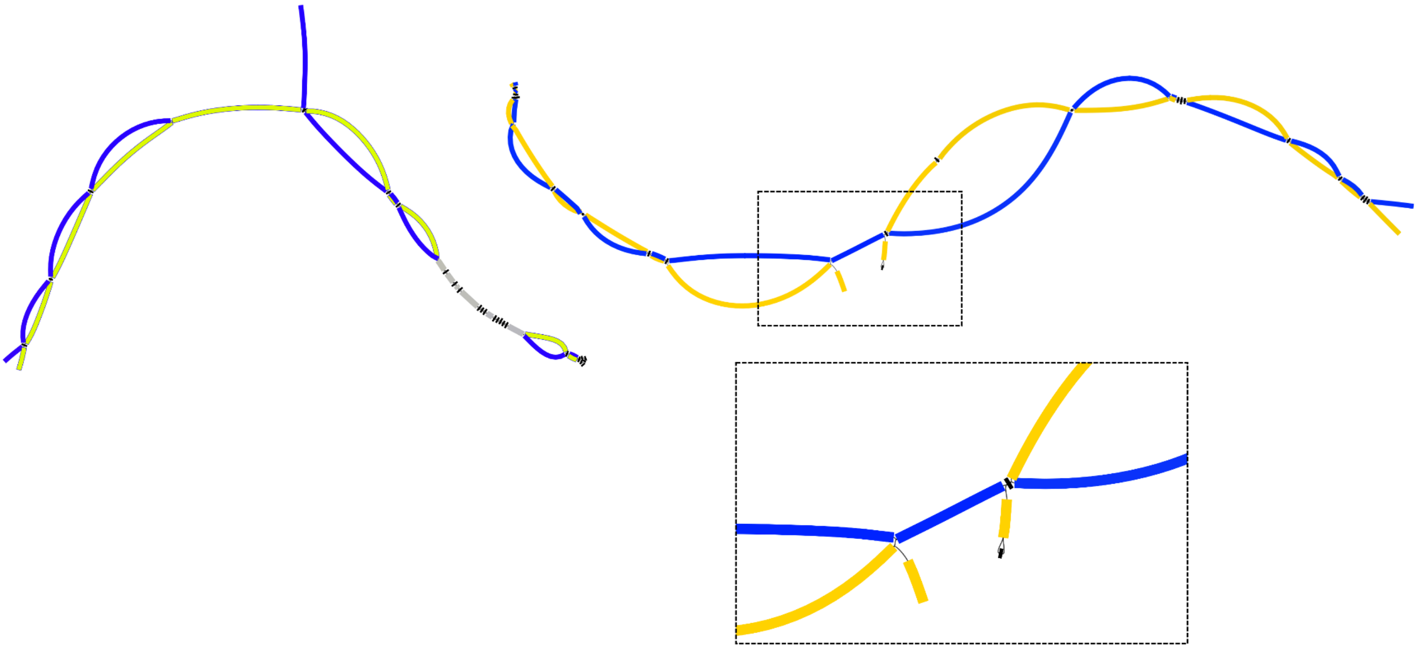

*Figure S2. Hairpin-Capped Broken Bubbles interrupt Rukki Scaffolding. Bandage [67] graphs of chromosomes 5 (left) and 1 (right, top) from the Verkko HG00733 assembly both show broken bubbles. On each connected component, the blue or yellow shade corresponds to a haplotype as inferred by Graphasing, grey corresponds to homozygous nodes, and black to nodes without inferred phasing signal. The blue haplotype on chromosome 5 is scaffolded in one path, but the yellow haplotype of chromosome 1 is represented with two scaffolds, one on*

*either side of the broken bubble, and is the only pictured haplotype not connected in a single path. Zooming in (right, bottom) shows that one end of the broken bubble is "capped" by a single small node in a hairpin structure.*

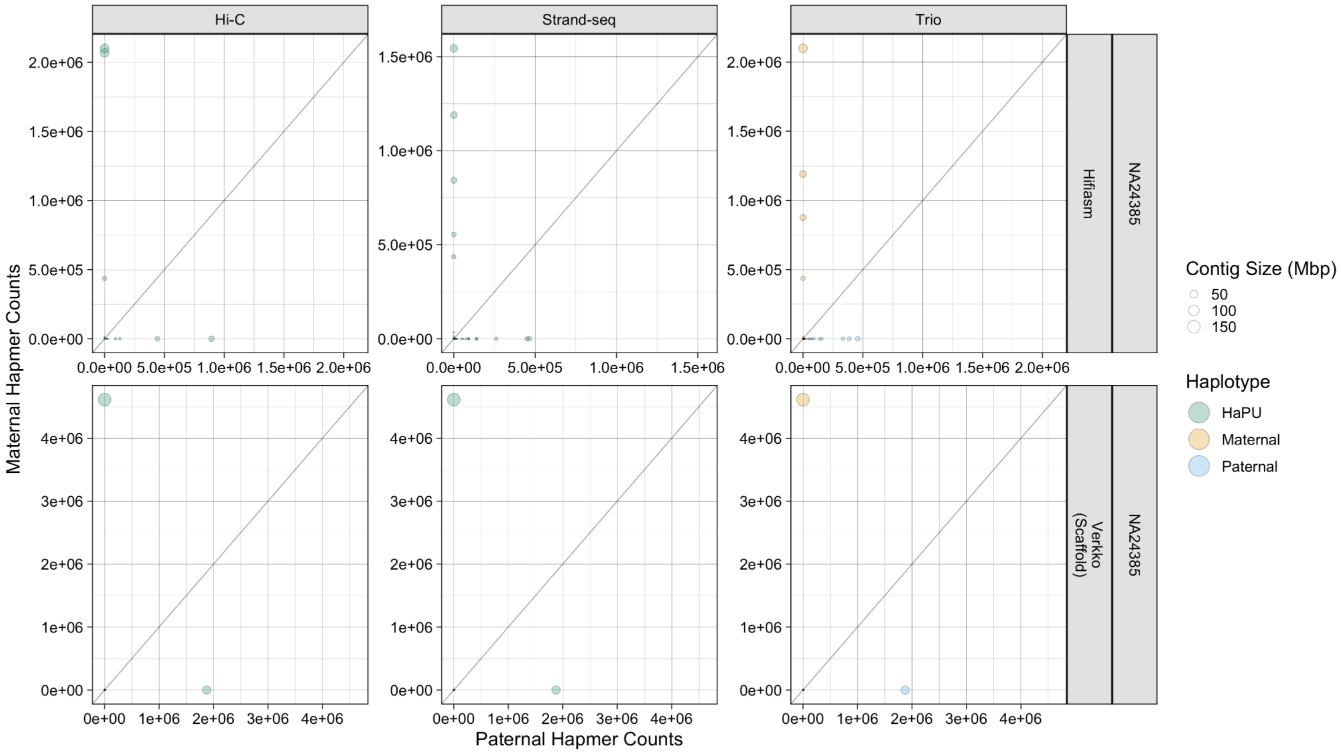


*Figure S3. Sex Chromosome Hap-mer Blob Plots. NA24385 contigs aligning to the X and Y chromosomes are plotted.*

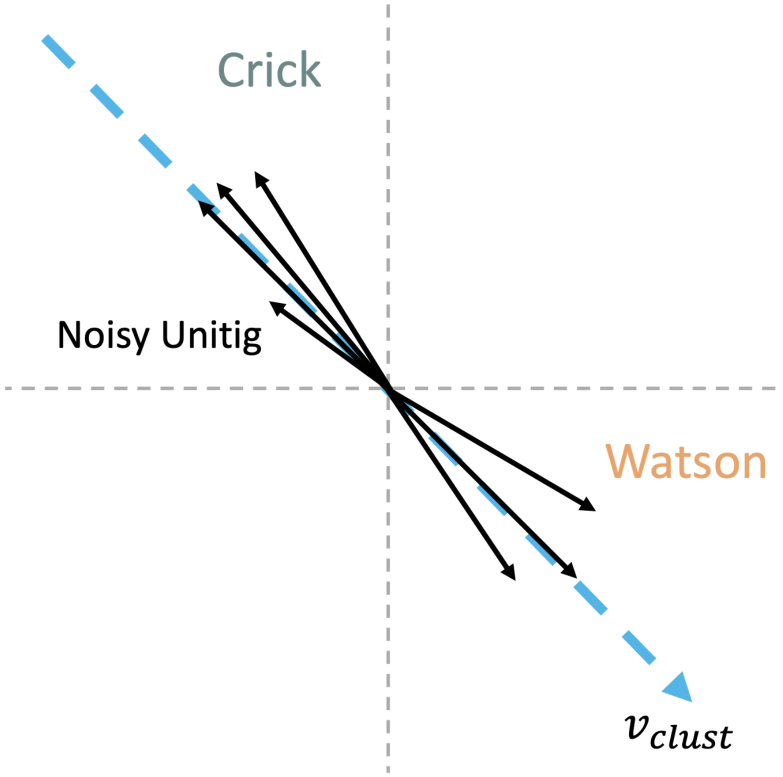


*Figure S4. Behavior of Absolute SSF Calculated with bwa mem alignments. Each axis corresponds to a different Strand-seq library. Unitigs from the same chromosome, but in opposite orientation will point in opposite directions along the same cluster vector (blue dashed vector), but are nonetheless clustered together when the absolute cosine similarity is used. A unitig that contains noisy, degenerate sequence is skewed and shrunk towards the 0 vector, but still points along its corresponding cluster vector.*

| | Hap 1 | Hap 2 |
|---|---|---|
| WC Library 1 | Watson | Crick |
| WC Library 2 | Crick | Watson |
| WC Library 3 | Crick | Watson |
| WC Library 4 | Watson | Crick |

| | Hap 1 | Hap 2 |
|---|---|---|
| WC Library 1 | Crick | Watson |
| WC Library 2 | Crick | Watson |
| WC Library 3 | Crick | Watson |
| WC Library 4 | Crick | Watson |

*Figure S5. Read Labeling for Proper Strand-seq Library Pooling. In each Strand-seq library, the labels (Watson, Crick) of the reads aligning to each haplotype depend on the orientation of the unitigs and the inherited Strand-state of the unmatched strand state library (WC vs. CW library). However, swapping the labels of reads does not affect the steps before library pooling, and so the labeling within each library can safely be treated as if it were arbitrary. Left: Each haplotype receives only Watson or only Crick alignments within each library. However, the pooled library assigns each haplotype a 50/50 ratio of Watson and Crick alignments and fails to reflect the behavior at the individual library level. Right: The labels of libraries 1 and 4 are swapped, and the pooled library now also assigns reads of only one label to each haplotype.*

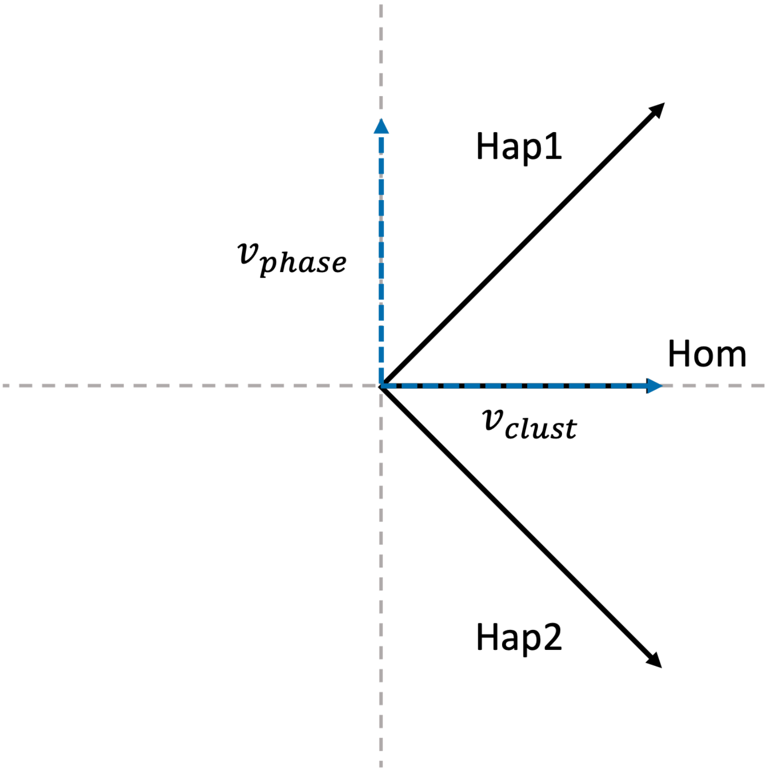


*Figure S6. Each axis corresponds to a different Strand-seq library. When the SSF is calculated with phase-informative reads, the unitig vectors (black arrows) separate from the cluster vector according to the haplotype. The difference between the haplotype 1 and haplotype 2 vectors defines the phase vector, which is used to pool Strand-seq libraries. The vectors of unitigs that contain homozygous sequence remain parallel to the cluster vector regardless of the reads used to calculate the SSF.*

Table S1: Unitig Filtering Statistics

| **Sample** | **Assembler** | **Total Unitigs (n)** | **Unitigs < 50kbp (n)** | **Total Assembly Size < 50kbp (%)** |
|---|---|---|---|---|
| NA24385 | Verkko hybrid | 8,627 | 4,966 | 2.2 |
| NA24385 | hifiasm hybrid | 12,520 | 4,355 | 2.5 |
| HG00733 | Verkko hybrid | 1,882 | 1,353 | 0.5 |
| HG00733 | hifiasm hybrid | 4,579 | 2,026 | 1.2 |

Table S2: NA24385 Strand-seq library quality annotations:

| **File 1** | **File 2** | **Annotation** |
|---|---|---|
| HWVTJAFXY_HG002x1xHG002x2_19s005136-1-1_Hasenfeld_lane1HG002x01PE20301_1_sequence.txt.gz | HWVTJAFXY_HG002x1xHG002x2_19s005136-1-1_Hasenfeld_lane1HG002x01PE20301_2_sequence.txt.gz | HighQuality |
| HWVTJAFXY_HG002x1xHG002x2_19s005136-1-1_Hasenfeld_lane1HG002x01PE20303_1_sequence.txt.gz | HWVTJAFXY_HG002x1xHG002x2_19s005136-1-1_Hasenfeld_lane1HG002x01PE20303_2_sequence.txt.gz | HighQuality |

| | | |
|---|---|---|
| HWVTJAFXY_HG002x1xHG002x2_19s005136-1-1_Hasenfeld_lane1HG002x01PE20305_1_sequence.txt.gz | HWVTJAFXY_HG002x1xHG002x2_19s005136-1-1_Hasenfeld_lane1HG002x01PE20305_2_sequence.txt.gz | HighQuality |
| HWVTJAFXY_HG002x1xHG002x2_19s005136-1-1_Hasenfeld_lane1HG002x01PE20306_1_sequence.txt.gz | HWVTJAFXY_HG002x1xHG002x2_19s005136-1-1_Hasenfeld_lane1HG002x01PE20306_2_sequence.txt.gz | HighQuality |
| HWVTJAFXY_HG002x1xHG002x2_19s005136-1-1_Hasenfeld_lane1HG002x01PE20307_1_sequence.txt.gz | HWVTJAFXY_HG002x1xHG002x2_19s005136-1-1_Hasenfeld_lane1HG002x01PE20307_2_sequence.txt.gz | HighQuality |
| HWVTJAFXY_HG002x1xHG002x2_19s005136-1-1_Hasenfeld_lane1HG002x01PE20308_1_sequence.txt.gz | HWVTJAFXY_HG002x1xHG002x2_19s005136-1-1_Hasenfeld_lane1HG002x01PE20308_2_sequence.txt.gz | HighQuality |
| HWVTJAFXY_HG002x1xHG002x2_19s005136-1-1_Hasenfeld_lane1HG002x01PE20313_1_sequence.txt.gz | HWVTJAFXY_HG002x1xHG002x2_19s005136-1-1_Hasenfeld_lane1HG002x01PE20313_2_sequence.txt.gz | HighQuality |
| HWVTJAFXY_HG002x1xHG002x2_19s005136-1-1_Hasenfeld_lane1HG002x01PE20315_1_sequence.txt.gz | HWVTJAFXY_HG002x1xHG002x2_19s005136-1-1_Hasenfeld_lane1HG002x01PE20315_2_sequence.txt.gz | HighQuality |
| HWVTJAFXY_HG002x1xHG002x2_19s005136-1-1_Hasenfeld_lane1HG002x01PE20318_1_sequence.txt.gz | HWVTJAFXY_HG002x1xHG002x2_19s005136-1-1_Hasenfeld_lane1HG002x01PE20318_2_sequence.txt.gz | HighQuality |
| HWVTJAFXY_HG002x1xHG002x2_19s005136-1-1_Hasenfeld_lane1HG002x01PE20319_1_sequence.txt.gz | HWVTJAFXY_HG002x1xHG002x2_19s005136-1-1_Hasenfeld_lane1HG002x01PE20319_2_sequence.txt.gz | HighQuality |
| HWVTJAFXY_HG002x1xHG002x2_19s005136-1-1_Hasenfeld_lane1HG002x01PE20325_1_sequence.txt.gz | HWVTJAFXY_HG002x1xHG002x2_19s005136-1-1_Hasenfeld_lane1HG002x01PE20325_2_sequence.txt.gz | HighQuality |
| HWVTJAFXY_HG002x1xHG002x2_19s005136-1-1_Hasenfeld_lane1HG002x01PE20327_1_sequence.txt.gz | HWVTJAFXY_HG002x1xHG002x2_19s005136-1-1_Hasenfeld_lane1HG002x01PE20327_2_sequence.txt.gz | HighQuality |
| HWVTJAFXY_HG002x1xHG002x2_19s005136-1-1_Hasenfeld_lane1HG002x01PE20328_1_sequence.txt.gz | HWVTJAFXY_HG002x1xHG002x2_19s005136-1-1_Hasenfeld_lane1HG002x01PE20328_2_sequence.txt.gz | HighQuality |
| HWVTJAFXY_HG002x1xHG002x2_19s005136-1-1_Hasenfeld_lane1HG002x01PE20329_1_sequence.txt.gz | HWVTJAFXY_HG002x1xHG002x2_19s005136-1-1_Hasenfeld_lane1HG002x01PE20329_2_sequence.txt.gz | HighQuality |
| HWVTJAFXY_HG002x1xHG002x2_19s005136-1-1_Hasenfeld_lane1HG002x01PE20331_1_sequence.txt.gz | HWVTJAFXY_HG002x1xHG002x2_19s005136-1-1_Hasenfeld_lane1HG002x01PE20331_2_sequence.txt.gz | HighQuality |
| HWVTJAFXY_HG002x1xHG002x2_19s005136-1-1_Hasenfeld_lane1HG002x01PE20332_1_sequence.txt.gz | HWVTJAFXY_HG002x1xHG002x2_19s005136-1-1_Hasenfeld_lane1HG002x01PE20332_2_sequence.txt.gz | HighQuality |
| HWVTJAFXY_HG002x1xHG002x2_19s005136-1-1_Hasenfeld_lane1HG002x01PE20334_1_sequence.txt.gz | HWVTJAFXY_HG002x1xHG002x2_19s005136-1-1_Hasenfeld_lane1HG002x01PE20334_2_sequence.txt.gz | HighQuality |
| HWVTJAFXY_HG002x1xHG002x2_19s005136-1-1_Hasenfeld_lane1HG002x01PE20335_1_sequence.txt.gz | HWVTJAFXY_HG002x1xHG002x2_19s005136-1-1_Hasenfeld_lane1HG002x01PE20335_2_sequence.txt.gz | HighQuality |
| HWVTJAFXY_HG002x1xHG002x2_19s005136-1-1_Hasenfeld_lane1HG002x01PE20336_1_sequence.txt.gz | HWVTJAFXY_HG002x1xHG002x2_19s005136-1-1_Hasenfeld_lane1HG002x01PE20336_2_sequence.txt.gz | HighQuality |
| HWVTJAFXY_HG002x1xHG002x2_19s005136-1-1_Hasenfeld_lane1HG002x01PE20337_1_sequence.txt.gz | HWVTJAFXY_HG002x1xHG002x2_19s005136-1-1_Hasenfeld_lane1HG002x01PE20337_2_sequence.txt.gz | HighQuality |
| HWVTJAFXY_HG002x1xHG002x2_19s005136-1-1_Hasenfeld_lane1HG002x01PE20339_1_sequence.txt.gz | HWVTJAFXY_HG002x1xHG002x2_19s005136-1-1_Hasenfeld_lane1HG002x01PE20339_2_sequence.txt.gz | HighQuality |
| HWVTJAFXY_HG002x1xHG002x2_19s005136-1-1_Hasenfeld_lane1HG002x01PE20340_1_sequence.txt.gz | HWVTJAFXY_HG002x1xHG002x2_19s005136-1-1_Hasenfeld_lane1HG002x01PE20340_2_sequence.txt.gz | HighQuality |
| HWVTJAFXY_HG002x1xHG002x2_19s005136-1-1_Hasenfeld_lane1HG002x01PE20341_1_sequence.txt.gz | HWVTJAFXY_HG002x1xHG002x2_19s005136-1-1_Hasenfeld_lane1HG002x01PE20341_2_sequence.txt.gz | HighQuality |
| HWVTJAFXY_HG002x1xHG002x2_19s005136-1-1_Hasenfeld_lane1HG002x01PE20342_1_sequence.txt.gz | HWVTJAFXY_HG002x1xHG002x2_19s005136-1-1_Hasenfeld_lane1HG002x01PE20342_2_sequence.txt.gz | HighQuality |
| HWVTJAFXY_HG002x1xHG002x2_19s005136-1-1_Hasenfeld_lane1HG002x01PE20343_1_sequence.txt.gz | HWVTJAFXY_HG002x1xHG002x2_19s005136-1-1_Hasenfeld_lane1HG002x01PE20343_2_sequence.txt.gz | HighQuality |
| HWVTJAFXY_HG002x1xHG002x2_19s005136-1-1_Hasenfeld_lane1HG002x01PE20345_1_sequence.txt.gz | HWVTJAFXY_HG002x1xHG002x2_19s005136-1-1_Hasenfeld_lane1HG002x01PE20345_2_sequence.txt.gz | HighQuality |
| HWVTJAFXY_HG002x1xHG002x2_19s005136-1-1_Hasenfeld_lane1HG002x01PE20347_1_sequence.txt.gz | HWVTJAFXY_HG002x1xHG002x2_19s005136-1-1_Hasenfeld_lane1HG002x01PE20347_2_sequence.txt.gz | HighQuality |

| | | |
|---|---|---|
| HWVTJAFXY_HG002x1xHG002x2_19s005136-1-1_Hasenfeld_lane1HG002x01PE20350_1_sequence.txt.gz | HWVTJAFXY_HG002x1xHG002x2_19s005136-1-1_Hasenfeld_lane1HG002x01PE20350_2_sequence.txt.gz | HighQuality |
| HWVTJAFXY_HG002x1xHG002x2_19s005136-1-1_Hasenfeld_lane1HG002x01PE20351_1_sequence.txt.gz | HWVTJAFXY_HG002x1xHG002x2_19s005136-1-1_Hasenfeld_lane1HG002x01PE20351_2_sequence.txt.gz | HighQuality |
| HWVTJAFXY_HG002x1xHG002x2_19s005136-1-1_Hasenfeld_lane1HG002x01PE20352_1_sequence.txt.gz | HWVTJAFXY_HG002x1xHG002x2_19s005136-1-1_Hasenfeld_lane1HG002x01PE20352_2_sequence.txt.gz | HighQuality |
| HWVTJAFXY_HG002x1xHG002x2_19s005136-1-1_Hasenfeld_lane1HG002x01PE20353_1_sequence.txt.gz | HWVTJAFXY_HG002x1xHG002x2_19s005136-1-1_Hasenfeld_lane1HG002x01PE20353_2_sequence.txt.gz | HighQuality |
| HWVTJAFXY_HG002x1xHG002x2_19s005136-1-1_Hasenfeld_lane1HG002x01PE20355_1_sequence.txt.gz | HWVTJAFXY_HG002x1xHG002x2_19s005136-1-1_Hasenfeld_lane1HG002x01PE20355_2_sequence.txt.gz | HighQuality |
| HWVTJAFXY_HG002x1xHG002x2_19s005136-1-1_Hasenfeld_lane1HG002x01PE20356_1_sequence.txt.gz | HWVTJAFXY_HG002x1xHG002x2_19s005136-1-1_Hasenfeld_lane1HG002x01PE20356_2_sequence.txt.gz | HighQuality |
| HWVTJAFXY_HG002x1xHG002x2_19s005136-1-1_Hasenfeld_lane1HG002x01PE20357_1_sequence.txt.gz | HWVTJAFXY_HG002x1xHG002x2_19s005136-1-1_Hasenfeld_lane1HG002x01PE20357_2_sequence.txt.gz | HighQuality |
| HWVTJAFXY_HG002x1xHG002x2_19s005136-1-1_Hasenfeld_lane1HG002x01PE20358_1_sequence.txt.gz | HWVTJAFXY_HG002x1xHG002x2_19s005136-1-1_Hasenfeld_lane1HG002x01PE20358_2_sequence.txt.gz | HighQuality |
| HWVTJAFXY_HG002x1xHG002x2_19s005136-1-1_Hasenfeld_lane1HG002x01PE20359_1_sequence.txt.gz | HWVTJAFXY_HG002x1xHG002x2_19s005136-1-1_Hasenfeld_lane1HG002x01PE20359_2_sequence.txt.gz | HighQuality |
| HWVTJAFXY_HG002x1xHG002x2_19s005136-1-1_Hasenfeld_lane1HG002x01PE20361_1_sequence.txt.gz | HWVTJAFXY_HG002x1xHG002x2_19s005136-1-1_Hasenfeld_lane1HG002x01PE20361_2_sequence.txt.gz | HighQuality |
| HWVTJAFXY_HG002x1xHG002x2_19s005136-1-1_Hasenfeld_lane1HG002x01PE20362_1_sequence.txt.gz | HWVTJAFXY_HG002x1xHG002x2_19s005136-1-1_Hasenfeld_lane1HG002x01PE20362_2_sequence.txt.gz | HighQuality |
| HWVTJAFXY_HG002x1xHG002x2_19s005136-1-1_Hasenfeld_lane1HG002x01PE20363_1_sequence.txt.gz | HWVTJAFXY_HG002x1xHG002x2_19s005136-1-1_Hasenfeld_lane1HG002x01PE20363_2_sequence.txt.gz | HighQuality |
| HWVTJAFXY_HG002x1xHG002x2_19s005136-1-1_Hasenfeld_lane1HG002x01PE20364_1_sequence.txt.gz | HWVTJAFXY_HG002x1xHG002x2_19s005136-1-1_Hasenfeld_lane1HG002x01PE20364_2_sequence.txt.gz | HighQuality |
| HWVTJAFXY_HG002x1xHG002x2_19s005136-1-1_Hasenfeld_lane1HG002x01PE20367_1_sequence.txt.gz | HWVTJAFXY_HG002x1xHG002x2_19s005136-1-1_Hasenfeld_lane1HG002x01PE20367_2_sequence.txt.gz | HighQuality |
| HWVTJAFXY_HG002x1xHG002x2_19s005136-1-1_Hasenfeld_lane1HG002x01PE20368_1_sequence.txt.gz | HWVTJAFXY_HG002x1xHG002x2_19s005136-1-1_Hasenfeld_lane1HG002x01PE20368_2_sequence.txt.gz | HighQuality |
| HWVTJAFXY_HG002x1xHG002x2_19s005136-1-1_Hasenfeld_lane1HG002x01PE20374_1_sequence.txt.gz | HWVTJAFXY_HG002x1xHG002x2_19s005136-1-1_Hasenfeld_lane1HG002x01PE20374_2_sequence.txt.gz | HighQuality |
| HWVTJAFXY_HG002x1xHG002x2_19s005136-1-1_Hasenfeld_lane1HG002x01PE20376_1_sequence.txt.gz | HWVTJAFXY_HG002x1xHG002x2_19s005136-1-1_Hasenfeld_lane1HG002x01PE20376_2_sequence.txt.gz | HighQuality |
| HWVTJAFXY_HG002x1xHG002x2_19s005136-1-1_Hasenfeld_lane1HG002x01PE20377_1_sequence.txt.gz | HWVTJAFXY_HG002x1xHG002x2_19s005136-1-1_Hasenfeld_lane1HG002x01PE20377_2_sequence.txt.gz | HighQuality |
| HWVTJAFXY_HG002x1xHG002x2_19s005136-1-1_Hasenfeld_lane1HG002x01PE20378_1_sequence.txt.gz | HWVTJAFXY_HG002x1xHG002x2_19s005136-1-1_Hasenfeld_lane1HG002x01PE20378_2_sequence.txt.gz | HighQuality |
| HWVTJAFXY_HG002x1xHG002x2_19s005136-1-1_Hasenfeld_lane1HG002x01PE20379_1_sequence.txt.gz | HWVTJAFXY_HG002x1xHG002x2_19s005136-1-1_Hasenfeld_lane1HG002x01PE20379_2_sequence.txt.gz | HighQuality |
| HWVTJAFXY_HG002x1xHG002x2_19s005136-1-1_Hasenfeld_lane1HG002x01PE20381_1_sequence.txt.gz | HWVTJAFXY_HG002x1xHG002x2_19s005136-1-1_Hasenfeld_lane1HG002x01PE20381_2_sequence.txt.gz | HighQuality |
| HWVTJAFXY_HG002x1xHG002x2_19s005136-1-1_Hasenfeld_lane1HG002x01PE20387_1_sequence.txt.gz | HWVTJAFXY_HG002x1xHG002x2_19s005136-1-1_Hasenfeld_lane1HG002x01PE20387_2_sequence.txt.gz | HighQuality |
| HWVTJAFXY_HG002x1xHG002x2_19s005136-1-1_Hasenfeld_lane1HG002x01PE20388_1_sequence.txt.gz | HWVTJAFXY_HG002x1xHG002x2_19s005136-1-1_Hasenfeld_lane1HG002x01PE20388_2_sequence.txt.gz | HighQuality |
| HWVTJAFXY_HG002x1xHG002x2_19s005136-1-1_Hasenfeld_lane1HG002x01PE20389_1_sequence.txt.gz | HWVTJAFXY_HG002x1xHG002x2_19s005136-1-1_Hasenfeld_lane1HG002x01PE20389_2_sequence.txt.gz | HighQuality |
| HWVTJAFXY_HG002x1xHG002x2_19s005136-1-1_Hasenfeld_lane1HG002x01PE20391_1_sequence.txt.gz | HWVTJAFXY_HG002x1xHG002x2_19s005136-1-1_Hasenfeld_lane1HG002x01PE20391_2_sequence.txt.gz | HighQuality |

| | | |
|---|---|---|
| HWVTJAFXY_HG002x1xHG002x2_19s005136-1-1_Hasenfeld_lane1HG002x01PE20392_1_sequence.txt.gz | HWVTJAFXY_HG002x1xHG002x2_19s005136-1-1_Hasenfeld_lane1HG002x01PE20392_2_sequence.txt.gz | HighQuality |
| HWVTJAFXY_HG002x1xHG002x2_19s005136-1-1_Hasenfeld_lane1HG002x01PE20393_1_sequence.txt.gz | HWVTJAFXY_HG002x1xHG002x2_19s005136-1-1_Hasenfeld_lane1HG002x01PE20393_2_sequence.txt.gz | HighQuality |
| HWVTJAFXY_HG002x1xHG002x2_19s005136-1-1_Hasenfeld_lane1HG002x02PE20403_1_sequence.txt.gz | HWVTJAFXY_HG002x1xHG002x2_19s005136-1-1_Hasenfeld_lane1HG002x02PE20403_2_sequence.txt.gz | HighQuality |
| HWVTJAFXY_HG002x1xHG002x2_19s005136-1-1_Hasenfeld_lane1HG002x02PE20405_1_sequence.txt.gz | HWVTJAFXY_HG002x1xHG002x2_19s005136-1-1_Hasenfeld_lane1HG002x02PE20405_2_sequence.txt.gz | HighQuality |
| HWVTJAFXY_HG002x1xHG002x2_19s005136-1-1_Hasenfeld_lane1HG002x02PE20407_1_sequence.txt.gz | HWVTJAFXY_HG002x1xHG002x2_19s005136-1-1_Hasenfeld_lane1HG002x02PE20407_2_sequence.txt.gz | HighQuality |
| HWVTJAFXY_HG002x1xHG002x2_19s005136-1-1_Hasenfeld_lane1HG002x02PE20414_1_sequence.txt.gz | HWVTJAFXY_HG002x1xHG002x2_19s005136-1-1_Hasenfeld_lane1HG002x02PE20414_2_sequence.txt.gz | HighQuality |
| HWVTJAFXY_HG002x1xHG002x2_19s005136-1-1_Hasenfeld_lane1HG002x02PE20416_1_sequence.txt.gz | HWVTJAFXY_HG002x1xHG002x2_19s005136-1-1_Hasenfeld_lane1HG002x02PE20416_2_sequence.txt.gz | HighQuality |
| HWVTJAFXY_HG002x1xHG002x2_19s005136-1-1_Hasenfeld_lane1HG002x02PE20417_1_sequence.txt.gz | HWVTJAFXY_HG002x1xHG002x2_19s005136-1-1_Hasenfeld_lane1HG002x02PE20417_2_sequence.txt.gz | HighQuality |
| HWVTJAFXY_HG002x1xHG002x2_19s005136-1-1_Hasenfeld_lane1HG002x02PE20419_1_sequence.txt.gz | HWVTJAFXY_HG002x1xHG002x2_19s005136-1-1_Hasenfeld_lane1HG002x02PE20419_2_sequence.txt.gz | HighQuality |
| HWVTJAFXY_HG002x1xHG002x2_19s005136-1-1_Hasenfeld_lane1HG002x02PE20421_1_sequence.txt.gz | HWVTJAFXY_HG002x1xHG002x2_19s005136-1-1_Hasenfeld_lane1HG002x02PE20421_2_sequence.txt.gz | HighQuality |
| HWVTJAFXY_HG002x1xHG002x2_19s005136-1-1_Hasenfeld_lane1HG002x02PE20422_1_sequence.txt.gz | HWVTJAFXY_HG002x1xHG002x2_19s005136-1-1_Hasenfeld_lane1HG002x02PE20422_2_sequence.txt.gz | HighQuality |
| HWVTJAFXY_HG002x1xHG002x2_19s005136-1-1_Hasenfeld_lane1HG002x02PE20424_1_sequence.txt.gz | HWVTJAFXY_HG002x1xHG002x2_19s005136-1-1_Hasenfeld_lane1HG002x02PE20424_2_sequence.txt.gz | HighQuality |
| HWVTJAFXY_HG002x1xHG002x2_19s005136-1-1_Hasenfeld_lane1HG002x02PE20425_1_sequence.txt.gz | HWVTJAFXY_HG002x1xHG002x2_19s005136-1-1_Hasenfeld_lane1HG002x02PE20425_2_sequence.txt.gz | HighQuality |
| HWVTJAFXY_HG002x1xHG002x2_19s005136-1-1_Hasenfeld_lane1HG002x02PE20428_1_sequence.txt.gz | HWVTJAFXY_HG002x1xHG002x2_19s005136-1-1_Hasenfeld_lane1HG002x02PE20428_2_sequence.txt.gz | HighQuality |
| HWVTJAFXY_HG002x1xHG002x2_19s005136-1-1_Hasenfeld_lane1HG002x02PE20430_1_sequence.txt.gz | HWVTJAFXY_HG002x1xHG002x2_19s005136-1-1_Hasenfeld_lane1HG002x02PE20430_2_sequence.txt.gz | HighQuality |
| HWVTJAFXY_HG002x1xHG002x2_19s005136-1-1_Hasenfeld_lane1HG002x02PE20431_1_sequence.txt.gz | HWVTJAFXY_HG002x1xHG002x2_19s005136-1-1_Hasenfeld_lane1HG002x02PE20431_2_sequence.txt.gz | HighQuality |
| HWVTJAFXY_HG002x1xHG002x2_19s005136-1-1_Hasenfeld_lane1HG002x02PE20432_1_sequence.txt.gz | HWVTJAFXY_HG002x1xHG002x2_19s005136-1-1_Hasenfeld_lane1HG002x02PE20432_2_sequence.txt.gz | HighQuality |
| HWVTJAFXY_HG002x1xHG002x2_19s005136-1-1_Hasenfeld_lane1HG002x02PE20434_1_sequence.txt.gz | HWVTJAFXY_HG002x1xHG002x2_19s005136-1-1_Hasenfeld_lane1HG002x02PE20434_2_sequence.txt.gz | HighQuality |
| HWVTJAFXY_HG002x1xHG002x2_19s005136-1-1_Hasenfeld_lane1HG002x02PE20435_1_sequence.txt.gz | HWVTJAFXY_HG002x1xHG002x2_19s005136-1-1_Hasenfeld_lane1HG002x02PE20435_2_sequence.txt.gz | HighQuality |
| HWVTJAFXY_HG002x1xHG002x2_19s005136-1-1_Hasenfeld_lane1HG002x02PE20439_1_sequence.txt.gz | HWVTJAFXY_HG002x1xHG002x2_19s005136-1-1_Hasenfeld_lane1HG002x02PE20439_2_sequence.txt.gz | HighQuality |
| HWVTJAFXY_HG002x1xHG002x2_19s005136-1-1_Hasenfeld_lane1HG002x02PE20440_1_sequence.txt.gz | HWVTJAFXY_HG002x1xHG002x2_19s005136-1-1_Hasenfeld_lane1HG002x02PE20440_2_sequence.txt.gz | HighQuality |
| HWVTJAFXY_HG002x1xHG002x2_19s005136-1-1_Hasenfeld_lane1HG002x02PE20443_1_sequence.txt.gz | HWVTJAFXY_HG002x1xHG002x2_19s005136-1-1_Hasenfeld_lane1HG002x02PE20443_2_sequence.txt.gz | HighQuality |
| HWVTJAFXY_HG002x1xHG002x2_19s005136-1-1_Hasenfeld_lane1HG002x02PE20445_1_sequence.txt.gz | HWVTJAFXY_HG002x1xHG002x2_19s005136-1-1_Hasenfeld_lane1HG002x02PE20445_2_sequence.txt.gz | HighQuality |
| HWVTJAFXY_HG002x1xHG002x2_19s005136-1-1_Hasenfeld_lane1HG002x02PE20450_1_sequence.txt.gz | HWVTJAFXY_HG002x1xHG002x2_19s005136-1-1_Hasenfeld_lane1HG002x02PE20450_2_sequence.txt.gz | HighQuality |
| HWVTJAFXY_HG002x1xHG002x2_19s005136-1-1_Hasenfeld_lane1HG002x02PE20452_1_sequence.txt.gz | HWVTJAFXY_HG002x1xHG002x2_19s005136-1-1_Hasenfeld_lane1HG002x02PE20452_2_sequence.txt.gz | HighQuality |

| | | |
|---|---|---|
| HWVTJAFXY_HG002x1xHG002x2_19s005136-1-1_Hasenfeld_lane1HG002x02PE20454_1_sequence.txt.gz | HWVTJAFXY_HG002x1xHG002x2_19s005136-1-1_Hasenfeld_lane1HG002x02PE20454_2_sequence.txt.gz | HighQuality |
| HWVTJAFXY_HG002x1xHG002x2_19s005136-1-1_Hasenfeld_lane1HG002x02PE20456_1_sequence.txt.gz | HWVTJAFXY_HG002x1xHG002x2_19s005136-1-1_Hasenfeld_lane1HG002x02PE20456_2_sequence.txt.gz | HighQuality |
| HWVTJAFXY_HG002x1xHG002x2_19s005136-1-1_Hasenfeld_lane1HG002x02PE20458_1_sequence.txt.gz | HWVTJAFXY_HG002x1xHG002x2_19s005136-1-1_Hasenfeld_lane1HG002x02PE20458_2_sequence.txt.gz | HighQuality |
| HWVTJAFXY_HG002x1xHG002x2_19s005136-1-1_Hasenfeld_lane1HG002x02PE20464_1_sequence.txt.gz | HWVTJAFXY_HG002x1xHG002x2_19s005136-1-1_Hasenfeld_lane1HG002x02PE20464_2_sequence.txt.gz | HighQuality |
| HWVTJAFXY_HG002x1xHG002x2_19s005136-1-1_Hasenfeld_lane1HG002x02PE20466_1_sequence.txt.gz | HWVTJAFXY_HG002x1xHG002x2_19s005136-1-1_Hasenfeld_lane1HG002x02PE20466_2_sequence.txt.gz | HighQuality |
| HWVTJAFXY_HG002x1xHG002x2_19s005136-1-1_Hasenfeld_lane1HG002x02PE20467_1_sequence.txt.gz | HWVTJAFXY_HG002x1xHG002x2_19s005136-1-1_Hasenfeld_lane1HG002x02PE20467_2_sequence.txt.gz | HighQuality |
| HWVTJAFXY_HG002x1xHG002x2_19s005136-1-1_Hasenfeld_lane1HG002x02PE20469_1_sequence.txt.gz | HWVTJAFXY_HG002x1xHG002x2_19s005136-1-1_Hasenfeld_lane1HG002x02PE20469_2_sequence.txt.gz | HighQuality |
| HWVTJAFXY_HG002x1xHG002x2_19s005136-1-1_Hasenfeld_lane1HG002x02PE20473_1_sequence.txt.gz | HWVTJAFXY_HG002x1xHG002x2_19s005136-1-1_Hasenfeld_lane1HG002x02PE20473_2_sequence.txt.gz | HighQuality |
| HWVTJAFXY_HG002x1xHG002x2_19s005136-1-1_Hasenfeld_lane1HG002x02PE20479_1_sequence.txt.gz | HWVTJAFXY_HG002x1xHG002x2_19s005136-1-1_Hasenfeld_lane1HG002x02PE20479_2_sequence.txt.gz | HighQuality |
| HWVTJAFXY_HG002x1xHG002x2_19s005136-1-1_Hasenfeld_lane1HG002x02PE20483_1_sequence.txt.gz | HWVTJAFXY_HG002x1xHG002x2_19s005136-1-1_Hasenfeld_lane1HG002x02PE20483_2_sequence.txt.gz | HighQuality |
| HWVTJAFXY_HG002x1xHG002x2_19s005136-1-1_Hasenfeld_lane1HG002x02PE20484_1_sequence.txt.gz | HWVTJAFXY_HG002x1xHG002x2_19s005136-1-1_Hasenfeld_lane1HG002x02PE20484_2_sequence.txt.gz | HighQuality |
| HWVTJAFXY_HG002x1xHG002x2_19s005136-1-1_Hasenfeld_lane1HG002x02PE20485_1_sequence.txt.gz | HWVTJAFXY_HG002x1xHG002x2_19s005136-1-1_Hasenfeld_lane1HG002x02PE20485_2_sequence.txt.gz | HighQuality |
| HWVTJAFXY_HG002x1xHG002x2_19s005136-1-1_Hasenfeld_lane1HG002x02PE20486_1_sequence.txt.gz | HWVTJAFXY_HG002x1xHG002x2_19s005136-1-1_Hasenfeld_lane1HG002x02PE20486_2_sequence.txt.gz | HighQuality |
| HWVTJAFXY_HG002x1xHG002x2_19s005136-1-1_Hasenfeld_lane1HG002x02PE20487_1_sequence.txt.gz | HWVTJAFXY_HG002x1xHG002x2_19s005136-1-1_Hasenfeld_lane1HG002x02PE20487_2_sequence.txt.gz | HighQuality |
| HWVTJAFXY_HG002x1xHG002x2_19s005136-1-1_Hasenfeld_lane1HG002x02PE20488_1_sequence.txt.gz | HWVTJAFXY_HG002x1xHG002x2_19s005136-1-1_Hasenfeld_lane1HG002x02PE20488_2_sequence.txt.gz | HighQuality |
| HWVTJAFXY_HG002x1xHG002x2_19s005136-1-1_Hasenfeld_lane1HG002x02PE20490_1_sequence.txt.gz | HWVTJAFXY_HG002x1xHG002x2_19s005136-1-1_Hasenfeld_lane1HG002x02PE20490_2_sequence.txt.gz | HighQuality |
| HWVTJAFXY_HG002x1xHG002x2_19s005136-1-1_Hasenfeld_lane1HG002x02PE20491_1_sequence.txt.gz | HWVTJAFXY_HG002x1xHG002x2_19s005136-1-1_Hasenfeld_lane1HG002x02PE20491_2_sequence.txt.gz | HighQuality |
| HWVTJAFXY_HG002x1xHG002x2_19s005136-1-1_Hasenfeld_lane1HG002x02PE20492_1_sequence.txt.gz | HWVTJAFXY_HG002x1xHG002x2_19s005136-1-1_Hasenfeld_lane1HG002x02PE20492_2_sequence.txt.gz | HighQuality |
| HWVTJAFXY_HG002x1xHG002x2_19s005136-1-1_Hasenfeld_lane1HG002x02PE20494_1_sequence.txt.gz | HWVTJAFXY_HG002x1xHG002x2_19s005136-1-1_Hasenfeld_lane1HG002x02PE20494_2_sequence.txt.gz | HighQuality |
| HWVTJAFXY_HG002x1xHG002x2_19s005136-1-1_Hasenfeld_lane1HG002x01PE20302_1_sequence.txt.gz | HWVTJAFXY_HG002x1xHG002x2_19s005136-1-1_Hasenfeld_lane1HG002x01PE20302_2_sequence.txt.gz | NotHighQuality |
| HWVTJAFXY_HG002x1xHG002x2_19s005136-1-1_Hasenfeld_lane1HG002x01PE20304_1_sequence.txt.gz | HWVTJAFXY_HG002x1xHG002x2_19s005136-1-1_Hasenfeld_lane1HG002x01PE20304_2_sequence.txt.gz | NotHighQuality |
| HWVTJAFXY_HG002x1xHG002x2_19s005136-1-1_Hasenfeld_lane1HG002x01PE20309_1_sequence.txt.gz | HWVTJAFXY_HG002x1xHG002x2_19s005136-1-1_Hasenfeld_lane1HG002x01PE20309_2_sequence.txt.gz | NotHighQuality |
| HWVTJAFXY_HG002x1xHG002x2_19s005136-1-1_Hasenfeld_lane1HG002x01PE20310_1_sequence.txt.gz | HWVTJAFXY_HG002x1xHG002x2_19s005136-1-1_Hasenfeld_lane1HG002x01PE20310_2_sequence.txt.gz | NotHighQuality |
| HWVTJAFXY_HG002x1xHG002x2_19s005136-1-1_Hasenfeld_lane1HG002x01PE20311_1_sequence.txt.gz | HWVTJAFXY_HG002x1xHG002x2_19s005136-1-1_Hasenfeld_lane1HG002x01PE20311_2_sequence.txt.gz | NotHighQuality |
| HWVTJAFXY_HG002x1xHG002x2_19s005136-1-1_Hasenfeld_lane1HG002x01PE20312_1_sequence.txt.gz | HWVTJAFXY_HG002x1xHG002x2_19s005136-1-1_Hasenfeld_lane1HG002x01PE20312_2_sequence.txt.gz | NotHighQuality |

| | | |
|---|---|---|
| HWVTJAFXY_HG002x1xHG002x2_19s005136-1-1_Hasenfeld_lane1HG002x01PE20314_1_sequence.txt.gz | HWVTJAFXY_HG002x1xHG002x2_19s005136-1-1_Hasenfeld_lane1HG002x01PE20314_2_sequence.txt.gz | NotHighQuality |
| HWVTJAFXY_HG002x1xHG002x2_19s005136-1-1_Hasenfeld_lane1HG002x01PE20316_1_sequence.txt.gz | HWVTJAFXY_HG002x1xHG002x2_19s005136-1-1_Hasenfeld_lane1HG002x01PE20316_2_sequence.txt.gz | NotHighQuality |
| HWVTJAFXY_HG002x1xHG002x2_19s005136-1-1_Hasenfeld_lane1HG002x01PE20317_1_sequence.txt.gz | HWVTJAFXY_HG002x1xHG002x2_19s005136-1-1_Hasenfeld_lane1HG002x01PE20317_2_sequence.txt.gz | NotHighQuality |
| HWVTJAFXY_HG002x1xHG002x2_19s005136-1-1_Hasenfeld_lane1HG002x01PE20320_1_sequence.txt.gz | HWVTJAFXY_HG002x1xHG002x2_19s005136-1-1_Hasenfeld_lane1HG002x01PE20320_2_sequence.txt.gz | NotHighQuality |
| HWVTJAFXY_HG002x1xHG002x2_19s005136-1-1_Hasenfeld_lane1HG002x01PE20321_1_sequence.txt.gz | HWVTJAFXY_HG002x1xHG002x2_19s005136-1-1_Hasenfeld_lane1HG002x01PE20321_2_sequence.txt.gz | NotHighQuality |
| HWVTJAFXY_HG002x1xHG002x2_19s005136-1-1_Hasenfeld_lane1HG002x01PE20322_1_sequence.txt.gz | HWVTJAFXY_HG002x1xHG002x2_19s005136-1-1_Hasenfeld_lane1HG002x01PE20322_2_sequence.txt.gz | NotHighQuality |
| HWVTJAFXY_HG002x1xHG002x2_19s005136-1-1_Hasenfeld_lane1HG002x01PE20323_1_sequence.txt.gz | HWVTJAFXY_HG002x1xHG002x2_19s005136-1-1_Hasenfeld_lane1HG002x01PE20323_2_sequence.txt.gz | NotHighQuality |
| HWVTJAFXY_HG002x1xHG002x2_19s005136-1-1_Hasenfeld_lane1HG002x01PE20324_1_sequence.txt.gz | HWVTJAFXY_HG002x1xHG002x2_19s005136-1-1_Hasenfeld_lane1HG002x01PE20324_2_sequence.txt.gz | NotHighQuality |
| HWVTJAFXY_HG002x1xHG002x2_19s005136-1-1_Hasenfeld_lane1HG002x01PE20326_1_sequence.txt.gz | HWVTJAFXY_HG002x1xHG002x2_19s005136-1-1_Hasenfeld_lane1HG002x01PE20326_2_sequence.txt.gz | NotHighQuality |
| HWVTJAFXY_HG002x1xHG002x2_19s005136-1-1_Hasenfeld_lane1HG002x01PE20330_1_sequence.txt.gz | HWVTJAFXY_HG002x1xHG002x2_19s005136-1-1_Hasenfeld_lane1HG002x01PE20330_2_sequence.txt.gz | NotHighQuality |
| HWVTJAFXY_HG002x1xHG002x2_19s005136-1-1_Hasenfeld_lane1HG002x01PE20333_1_sequence.txt.gz | HWVTJAFXY_HG002x1xHG002x2_19s005136-1-1_Hasenfeld_lane1HG002x01PE20333_2_sequence.txt.gz | NotHighQuality |
| HWVTJAFXY_HG002x1xHG002x2_19s005136-1-1_Hasenfeld_lane1HG002x01PE20338_1_sequence.txt.gz | HWVTJAFXY_HG002x1xHG002x2_19s005136-1-1_Hasenfeld_lane1HG002x01PE20338_2_sequence.txt.gz | NotHighQuality |
| HWVTJAFXY_HG002x1xHG002x2_19s005136-1-1_Hasenfeld_lane1HG002x01PE20344_1_sequence.txt.gz | HWVTJAFXY_HG002x1xHG002x2_19s005136-1-1_Hasenfeld_lane1HG002x01PE20344_2_sequence.txt.gz | NotHighQuality |
| HWVTJAFXY_HG002x1xHG002x2_19s005136-1-1_Hasenfeld_lane1HG002x01PE20346_1_sequence.txt.gz | HWVTJAFXY_HG002x1xHG002x2_19s005136-1-1_Hasenfeld_lane1HG002x01PE20346_2_sequence.txt.gz | NotHighQuality |
| HWVTJAFXY_HG002x1xHG002x2_19s005136-1-1_Hasenfeld_lane1HG002x01PE20348_1_sequence.txt.gz | HWVTJAFXY_HG002x1xHG002x2_19s005136-1-1_Hasenfeld_lane1HG002x01PE20348_2_sequence.txt.gz | NotHighQuality |
| HWVTJAFXY_HG002x1xHG002x2_19s005136-1-1_Hasenfeld_lane1HG002x01PE20349_1_sequence.txt.gz | HWVTJAFXY_HG002x1xHG002x2_19s005136-1-1_Hasenfeld_lane1HG002x01PE20349_2_sequence.txt.gz | NotHighQuality |
| HWVTJAFXY_HG002x1xHG002x2_19s005136-1-1_Hasenfeld_lane1HG002x01PE20354_1_sequence.txt.gz | HWVTJAFXY_HG002x1xHG002x2_19s005136-1-1_Hasenfeld_lane1HG002x01PE20354_2_sequence.txt.gz | NotHighQuality |
| HWVTJAFXY_HG002x1xHG002x2_19s005136-1-1_Hasenfeld_lane1HG002x01PE20360_1_sequence.txt.gz | HWVTJAFXY_HG002x1xHG002x2_19s005136-1-1_Hasenfeld_lane1HG002x01PE20360_2_sequence.txt.gz | NotHighQuality |
| HWVTJAFXY_HG002x1xHG002x2_19s005136-1-1_Hasenfeld_lane1HG002x01PE20365_1_sequence.txt.gz | HWVTJAFXY_HG002x1xHG002x2_19s005136-1-1_Hasenfeld_lane1HG002x01PE20365_2_sequence.txt.gz | NotHighQuality |
| HWVTJAFXY_HG002x1xHG002x2_19s005136-1-1_Hasenfeld_lane1HG002x01PE20366_1_sequence.txt.gz | HWVTJAFXY_HG002x1xHG002x2_19s005136-1-1_Hasenfeld_lane1HG002x01PE20366_2_sequence.txt.gz | NotHighQuality |
| HWVTJAFXY_HG002x1xHG002x2_19s005136-1-1_Hasenfeld_lane1HG002x01PE20369_1_sequence.txt.gz | HWVTJAFXY_HG002x1xHG002x2_19s005136-1-1_Hasenfeld_lane1HG002x01PE20369_2_sequence.txt.gz | NotHighQuality |
| HWVTJAFXY_HG002x1xHG002x2_19s005136-1-1_Hasenfeld_lane1HG002x01PE20370_1_sequence.txt.gz | HWVTJAFXY_HG002x1xHG002x2_19s005136-1-1_Hasenfeld_lane1HG002x01PE20370_2_sequence.txt.gz | NotHighQuality |
| HWVTJAFXY_HG002x1xHG002x2_19s005136-1-1_Hasenfeld_lane1HG002x01PE20371_1_sequence.txt.gz | HWVTJAFXY_HG002x1xHG002x2_19s005136-1-1_Hasenfeld_lane1HG002x01PE20371_2_sequence.txt.gz | NotHighQuality |
| HWVTJAFXY_HG002x1xHG002x2_19s005136-1-1_Hasenfeld_lane1HG002x01PE20372_1_sequence.txt.gz | HWVTJAFXY_HG002x1xHG002x2_19s005136-1-1_Hasenfeld_lane1HG002x01PE20372_2_sequence.txt.gz | NotHighQuality |
| HWVTJAFXY_HG002x1xHG002x2_19s005136-1-1_Hasenfeld_lane1HG002x01PE20373_1_sequence.txt.gz | HWVTJAFXY_HG002x1xHG002x2_19s005136-1-1_Hasenfeld_lane1HG002x01PE20373_2_sequence.txt.gz | NotHighQuality |

| | | |
|---|---|---|
| HWVTJAFXY_HG002x1xHG002x2_19s005136-1-1_Hasenfeld_lane1HG002x01PE20375_1_sequence.txt.gz | HWVTJAFXY_HG002x1xHG002x2_19s005136-1-1_Hasenfeld_lane1HG002x01PE20375_2_sequence.txt.gz | NotHighQuality |
| HWVTJAFXY_HG002x1xHG002x2_19s005136-1-1_Hasenfeld_lane1HG002x01PE20380_1_sequence.txt.gz | HWVTJAFXY_HG002x1xHG002x2_19s005136-1-1_Hasenfeld_lane1HG002x01PE20380_2_sequence.txt.gz | NotHighQuality |
| HWVTJAFXY_HG002x1xHG002x2_19s005136-1-1_Hasenfeld_lane1HG002x01PE20382_1_sequence.txt.gz | HWVTJAFXY_HG002x1xHG002x2_19s005136-1-1_Hasenfeld_lane1HG002x01PE20382_2_sequence.txt.gz | NotHighQuality |
| HWVTJAFXY_HG002x1xHG002x2_19s005136-1-1_Hasenfeld_lane1HG002x01PE20383_1_sequence.txt.gz | HWVTJAFXY_HG002x1xHG002x2_19s005136-1-1_Hasenfeld_lane1HG002x01PE20383_2_sequence.txt.gz | NotHighQuality |
| HWVTJAFXY_HG002x1xHG002x2_19s005136-1-1_Hasenfeld_lane1HG002x01PE20384_1_sequence.txt.gz | HWVTJAFXY_HG002x1xHG002x2_19s005136-1-1_Hasenfeld_lane1HG002x01PE20384_2_sequence.txt.gz | NotHighQuality |
| HWVTJAFXY_HG002x1xHG002x2_19s005136-1-1_Hasenfeld_lane1HG002x01PE20385_1_sequence.txt.gz | HWVTJAFXY_HG002x1xHG002x2_19s005136-1-1_Hasenfeld_lane1HG002x01PE20385_2_sequence.txt.gz | NotHighQuality |
| HWVTJAFXY_HG002x1xHG002x2_19s005136-1-1_Hasenfeld_lane1HG002x01PE20386_1_sequence.txt.gz | HWVTJAFXY_HG002x1xHG002x2_19s005136-1-1_Hasenfeld_lane1HG002x01PE20386_2_sequence.txt.gz | NotHighQuality |
| HWVTJAFXY_HG002x1xHG002x2_19s005136-1-1_Hasenfeld_lane1HG002x01PE20390_1_sequence.txt.gz | HWVTJAFXY_HG002x1xHG002x2_19s005136-1-1_Hasenfeld_lane1HG002x01PE20390_2_sequence.txt.gz | NotHighQuality |
| HWVTJAFXY_HG002x1xHG002x2_19s005136-1-1_Hasenfeld_lane1HG002x01PE20394_1_sequence.txt.gz | HWVTJAFXY_HG002x1xHG002x2_19s005136-1-1_Hasenfeld_lane1HG002x01PE20394_2_sequence.txt.gz | NotHighQuality |
| HWVTJAFXY_HG002x1xHG002x2_19s005136-1-1_Hasenfeld_lane1HG002x01PE20395_1_sequence.txt.gz | HWVTJAFXY_HG002x1xHG002x2_19s005136-1-1_Hasenfeld_lane1HG002x01PE20395_2_sequence.txt.gz | NotHighQuality |
| HWVTJAFXY_HG002x1xHG002x2_19s005136-1-1_Hasenfeld_lane1HG002x01PE20396_1_sequence.txt.gz | HWVTJAFXY_HG002x1xHG002x2_19s005136-1-1_Hasenfeld_lane1HG002x01PE20396_2_sequence.txt.gz | NotHighQuality |
| HWVTJAFXY_HG002x1xHG002x2_19s005136-1-1_Hasenfeld_lane1HG002x02PE20401_1_sequence.txt.gz | HWVTJAFXY_HG002x1xHG002x2_19s005136-1-1_Hasenfeld_lane1HG002x02PE20401_2_sequence.txt.gz | NotHighQuality |
| HWVTJAFXY_HG002x1xHG002x2_19s005136-1-1_Hasenfeld_lane1HG002x02PE20402_1_sequence.txt.gz | HWVTJAFXY_HG002x1xHG002x2_19s005136-1-1_Hasenfeld_lane1HG002x02PE20402_2_sequence.txt.gz | NotHighQuality |
| HWVTJAFXY_HG002x1xHG002x2_19s005136-1-1_Hasenfeld_lane1HG002x02PE20404_1_sequence.txt.gz | HWVTJAFXY_HG002x1xHG002x2_19s005136-1-1_Hasenfeld_lane1HG002x02PE20404_2_sequence.txt.gz | NotHighQuality |
| HWVTJAFXY_HG002x1xHG002x2_19s005136-1-1_Hasenfeld_lane1HG002x02PE20406_1_sequence.txt.gz | HWVTJAFXY_HG002x1xHG002x2_19s005136-1-1_Hasenfeld_lane1HG002x02PE20406_2_sequence.txt.gz | NotHighQuality |
| HWVTJAFXY_HG002x1xHG002x2_19s005136-1-1_Hasenfeld_lane1HG002x02PE20408_1_sequence.txt.gz | HWVTJAFXY_HG002x1xHG002x2_19s005136-1-1_Hasenfeld_lane1HG002x02PE20408_2_sequence.txt.gz | NotHighQuality |
| HWVTJAFXY_HG002x1xHG002x2_19s005136-1-1_Hasenfeld_lane1HG002x02PE20409_1_sequence.txt.gz | HWVTJAFXY_HG002x1xHG002x2_19s005136-1-1_Hasenfeld_lane1HG002x02PE20409_2_sequence.txt.gz | NotHighQuality |
| HWVTJAFXY_HG002x1xHG002x2_19s005136-1-1_Hasenfeld_lane1HG002x02PE20410_1_sequence.txt.gz | HWVTJAFXY_HG002x1xHG002x2_19s005136-1-1_Hasenfeld_lane1HG002x02PE20410_2_sequence.txt.gz | NotHighQuality |
| HWVTJAFXY_HG002x1xHG002x2_19s005136-1-1_Hasenfeld_lane1HG002x02PE20411_1_sequence.txt.gz | HWVTJAFXY_HG002x1xHG002x2_19s005136-1-1_Hasenfeld_lane1HG002x02PE20411_2_sequence.txt.gz | NotHighQuality |
| HWVTJAFXY_HG002x1xHG002x2_19s005136-1-1_Hasenfeld_lane1HG002x02PE20412_1_sequence.txt.gz | HWVTJAFXY_HG002x1xHG002x2_19s005136-1-1_Hasenfeld_lane1HG002x02PE20412_2_sequence.txt.gz | NotHighQuality |
| HWVTJAFXY_HG002x1xHG002x2_19s005136-1-1_Hasenfeld_lane1HG002x02PE20413_1_sequence.txt.gz | HWVTJAFXY_HG002x1xHG002x2_19s005136-1-1_Hasenfeld_lane1HG002x02PE20413_2_sequence.txt.gz | NotHighQuality |
| HWVTJAFXY_HG002x1xHG002x2_19s005136-1-1_Hasenfeld_lane1HG002x02PE20415_1_sequence.txt.gz | HWVTJAFXY_HG002x1xHG002x2_19s005136-1-1_Hasenfeld_lane1HG002x02PE20415_2_sequence.txt.gz | NotHighQuality |
| HWVTJAFXY_HG002x1xHG002x2_19s005136-1-1_Hasenfeld_lane1HG002x02PE20418_1_sequence.txt.gz | HWVTJAFXY_HG002x1xHG002x2_19s005136-1-1_Hasenfeld_lane1HG002x02PE20418_2_sequence.txt.gz | NotHighQuality |
| HWVTJAFXY_HG002x1xHG002x2_19s005136-1-1_Hasenfeld_lane1HG002x02PE20420_1_sequence.txt.gz | HWVTJAFXY_HG002x1xHG002x2_19s005136-1-1_Hasenfeld_lane1HG002x02PE20420_2_sequence.txt.gz | NotHighQuality |
| HWVTJAFXY_HG002x1xHG002x2_19s005136-1-1_Hasenfeld_lane1HG002x02PE20423_1_sequence.txt.gz | HWVTJAFXY_HG002x1xHG002x2_19s005136-1-1_Hasenfeld_lane1HG002x02PE20423_2_sequence.txt.gz | NotHighQuality |

| | | |
|---|---|---|
| HWVTJAFXY_HG002x1xHG002x2_19s005136-1-1_Hasenfeld_lane1HG002x02PE20426_1_sequence.txt.gz | HWVTJAFXY_HG002x1xHG002x2_19s005136-1-1_Hasenfeld_lane1HG002x02PE20426_2_sequence.txt.gz | NotHighQuality |
| HWVTJAFXY_HG002x1xHG002x2_19s005136-1-1_Hasenfeld_lane1HG002x02PE20427_1_sequence.txt.gz | HWVTJAFXY_HG002x1xHG002x2_19s005136-1-1_Hasenfeld_lane1HG002x02PE20427_2_sequence.txt.gz | NotHighQuality |
| HWVTJAFXY_HG002x1xHG002x2_19s005136-1-1_Hasenfeld_lane1HG002x02PE20429_1_sequence.txt.gz | HWVTJAFXY_HG002x1xHG002x2_19s005136-1-1_Hasenfeld_lane1HG002x02PE20429_2_sequence.txt.gz | NotHighQuality |
| HWVTJAFXY_HG002x1xHG002x2_19s005136-1-1_Hasenfeld_lane1HG002x02PE20433_1_sequence.txt.gz | HWVTJAFXY_HG002x1xHG002x2_19s005136-1-1_Hasenfeld_lane1HG002x02PE20433_2_sequence.txt.gz | NotHighQuality |
| HWVTJAFXY_HG002x1xHG002x2_19s005136-1-1_Hasenfeld_lane1HG002x02PE20436_1_sequence.txt.gz | HWVTJAFXY_HG002x1xHG002x2_19s005136-1-1_Hasenfeld_lane1HG002x02PE20436_2_sequence.txt.gz | NotHighQuality |
| HWVTJAFXY_HG002x1xHG002x2_19s005136-1-1_Hasenfeld_lane1HG002x02PE20437_1_sequence.txt.gz | HWVTJAFXY_HG002x1xHG002x2_19s005136-1-1_Hasenfeld_lane1HG002x02PE20437_2_sequence.txt.gz | NotHighQuality |
| HWVTJAFXY_HG002x1xHG002x2_19s005136-1-1_Hasenfeld_lane1HG002x02PE20438_1_sequence.txt.gz | HWVTJAFXY_HG002x1xHG002x2_19s005136-1-1_Hasenfeld_lane1HG002x02PE20438_2_sequence.txt.gz | NotHighQuality |
| HWVTJAFXY_HG002x1xHG002x2_19s005136-1-1_Hasenfeld_lane1HG002x02PE20441_1_sequence.txt.gz | HWVTJAFXY_HG002x1xHG002x2_19s005136-1-1_Hasenfeld_lane1HG002x02PE20441_2_sequence.txt.gz | NotHighQuality |
| HWVTJAFXY_HG002x1xHG002x2_19s005136-1-1_Hasenfeld_lane1HG002x02PE20442_1_sequence.txt.gz | HWVTJAFXY_HG002x1xHG002x2_19s005136-1-1_Hasenfeld_lane1HG002x02PE20442_2_sequence.txt.gz | NotHighQuality |
| HWVTJAFXY_HG002x1xHG002x2_19s005136-1-1_Hasenfeld_lane1HG002x02PE20444_1_sequence.txt.gz | HWVTJAFXY_HG002x1xHG002x2_19s005136-1-1_Hasenfeld_lane1HG002x02PE20444_2_sequence.txt.gz | NotHighQuality |
| HWVTJAFXY_HG002x1xHG002x2_19s005136-1-1_Hasenfeld_lane1HG002x02PE20446_1_sequence.txt.gz | HWVTJAFXY_HG002x1xHG002x2_19s005136-1-1_Hasenfeld_lane1HG002x02PE20446_2_sequence.txt.gz | NotHighQuality |
| HWVTJAFXY_HG002x1xHG002x2_19s005136-1-1_Hasenfeld_lane1HG002x02PE20447_1_sequence.txt.gz | HWVTJAFXY_HG002x1xHG002x2_19s005136-1-1_Hasenfeld_lane1HG002x02PE20447_2_sequence.txt.gz | NotHighQuality |
| HWVTJAFXY_HG002x1xHG002x2_19s005136-1-1_Hasenfeld_lane1HG002x02PE20448_1_sequence.txt.gz | HWVTJAFXY_HG002x1xHG002x2_19s005136-1-1_Hasenfeld_lane1HG002x02PE20448_2_sequence.txt.gz | NotHighQuality |
| HWVTJAFXY_HG002x1xHG002x2_19s005136-1-1_Hasenfeld_lane1HG002x02PE20449_1_sequence.txt.gz | HWVTJAFXY_HG002x1xHG002x2_19s005136-1-1_Hasenfeld_lane1HG002x02PE20449_2_sequence.txt.gz | NotHighQuality |
| HWVTJAFXY_HG002x1xHG002x2_19s005136-1-1_Hasenfeld_lane1HG002x02PE20451_1_sequence.txt.gz | HWVTJAFXY_HG002x1xHG002x2_19s005136-1-1_Hasenfeld_lane1HG002x02PE20451_2_sequence.txt.gz | NotHighQuality |
| HWVTJAFXY_HG002x1xHG002x2_19s005136-1-1_Hasenfeld_lane1HG002x02PE20453_1_sequence.txt.gz | HWVTJAFXY_HG002x1xHG002x2_19s005136-1-1_Hasenfeld_lane1HG002x02PE20453_2_sequence.txt.gz | NotHighQuality |
| HWVTJAFXY_HG002x1xHG002x2_19s005136-1-1_Hasenfeld_lane1HG002x02PE20455_1_sequence.txt.gz | HWVTJAFXY_HG002x1xHG002x2_19s005136-1-1_Hasenfeld_lane1HG002x02PE20455_2_sequence.txt.gz | NotHighQuality |
| HWVTJAFXY_HG002x1xHG002x2_19s005136-1-1_Hasenfeld_lane1HG002x02PE20457_1_sequence.txt.gz | HWVTJAFXY_HG002x1xHG002x2_19s005136-1-1_Hasenfeld_lane1HG002x02PE20457_2_sequence.txt.gz | NotHighQuality |
| HWVTJAFXY_HG002x1xHG002x2_19s005136-1-1_Hasenfeld_lane1HG002x02PE20459_1_sequence.txt.gz | HWVTJAFXY_HG002x1xHG002x2_19s005136-1-1_Hasenfeld_lane1HG002x02PE20459_2_sequence.txt.gz | NotHighQuality |
| HWVTJAFXY_HG002x1xHG002x2_19s005136-1-1_Hasenfeld_lane1HG002x02PE20460_1_sequence.txt.gz | HWVTJAFXY_HG002x1xHG002x2_19s005136-1-1_Hasenfeld_lane1HG002x02PE20460_2_sequence.txt.gz | NotHighQuality |
| HWVTJAFXY_HG002x1xHG002x2_19s005136-1-1_Hasenfeld_lane1HG002x02PE20461_1_sequence.txt.gz | HWVTJAFXY_HG002x1xHG002x2_19s005136-1-1_Hasenfeld_lane1HG002x02PE20461_2_sequence.txt.gz | NotHighQuality |
| HWVTJAFXY_HG002x1xHG002x2_19s005136-1-1_Hasenfeld_lane1HG002x02PE20462_1_sequence.txt.gz | HWVTJAFXY_HG002x1xHG002x2_19s005136-1-1_Hasenfeld_lane1HG002x02PE20462_2_sequence.txt.gz | NotHighQuality |
| HWVTJAFXY_HG002x1xHG002x2_19s005136-1-1_Hasenfeld_lane1HG002x02PE20463_1_sequence.txt.gz | HWVTJAFXY_HG002x1xHG002x2_19s005136-1-1_Hasenfeld_lane1HG002x02PE20463_2_sequence.txt.gz | NotHighQuality |
| HWVTJAFXY_HG002x1xHG002x2_19s005136-1-1_Hasenfeld_lane1HG002x02PE20465_1_sequence.txt.gz | HWVTJAFXY_HG002x1xHG002x2_19s005136-1-1_Hasenfeld_lane1HG002x02PE20465_2_sequence.txt.gz | NotHighQuality |
| HWVTJAFXY_HG002x1xHG002x2_19s005136-1-1_Hasenfeld_lane1HG002x02PE20468_1_sequence.txt.gz | HWVTJAFXY_HG002x1xHG002x2_19s005136-1-1_Hasenfeld_lane1HG002x02PE20468_2_sequence.txt.gz | NotHighQuality |

| | | |
|---|---|---|
| HWVTJAFXY_HG002x1xHG002x2_19s005136-1-1_Hasenfeld_lane1HG002x02PE20470_1_sequence.txt.gz | HWVTJAFXY_HG002x1xHG002x2_19s005136-1-1_Hasenfeld_lane1HG002x02PE20470_2_sequence.txt.gz | NotHighQuality |
| HWVTJAFXY_HG002x1xHG002x2_19s005136-1-1_Hasenfeld_lane1HG002x02PE20471_1_sequence.txt.gz | HWVTJAFXY_HG002x1xHG002x2_19s005136-1-1_Hasenfeld_lane1HG002x02PE20471_2_sequence.txt.gz | NotHighQuality |
| HWVTJAFXY_HG002x1xHG002x2_19s005136-1-1_Hasenfeld_lane1HG002x02PE20472_1_sequence.txt.gz | HWVTJAFXY_HG002x1xHG002x2_19s005136-1-1_Hasenfeld_lane1HG002x02PE20472_2_sequence.txt.gz | NotHighQuality |
| HWVTJAFXY_HG002x1xHG002x2_19s005136-1-1_Hasenfeld_lane1HG002x02PE20474_1_sequence.txt.gz | HWVTJAFXY_HG002x1xHG002x2_19s005136-1-1_Hasenfeld_lane1HG002x02PE20474_2_sequence.txt.gz | NotHighQuality |
| HWVTJAFXY_HG002x1xHG002x2_19s005136-1-1_Hasenfeld_lane1HG002x02PE20475_1_sequence.txt.gz | HWVTJAFXY_HG002x1xHG002x2_19s005136-1-1_Hasenfeld_lane1HG002x02PE20475_2_sequence.txt.gz | NotHighQuality |
| HWVTJAFXY_HG002x1xHG002x2_19s005136-1-1_Hasenfeld_lane1HG002x02PE20476_1_sequence.txt.gz | HWVTJAFXY_HG002x1xHG002x2_19s005136-1-1_Hasenfeld_lane1HG002x02PE20476_2_sequence.txt.gz | NotHighQuality |
| HWVTJAFXY_HG002x1xHG002x2_19s005136-1-1_Hasenfeld_lane1HG002x02PE20477_1_sequence.txt.gz | HWVTJAFXY_HG002x1xHG002x2_19s005136-1-1_Hasenfeld_lane1HG002x02PE20477_2_sequence.txt.gz | NotHighQuality |
| HWVTJAFXY_HG002x1xHG002x2_19s005136-1-1_Hasenfeld_lane1HG002x02PE20478_1_sequence.txt.gz | HWVTJAFXY_HG002x1xHG002x2_19s005136-1-1_Hasenfeld_lane1HG002x02PE20478_2_sequence.txt.gz | NotHighQuality |
| HWVTJAFXY_HG002x1xHG002x2_19s005136-1-1_Hasenfeld_lane1HG002x02PE20480_1_sequence.txt.gz | HWVTJAFXY_HG002x1xHG002x2_19s005136-1-1_Hasenfeld_lane1HG002x02PE20480_2_sequence.txt.gz | NotHighQuality |
| HWVTJAFXY_HG002x1xHG002x2_19s005136-1-1_Hasenfeld_lane1HG002x02PE20481_1_sequence.txt.gz | HWVTJAFXY_HG002x1xHG002x2_19s005136-1-1_Hasenfeld_lane1HG002x02PE20481_2_sequence.txt.gz | NotHighQuality |
| HWVTJAFXY_HG002x1xHG002x2_19s005136-1-1_Hasenfeld_lane1HG002x02PE20482_1_sequence.txt.gz | HWVTJAFXY_HG002x1xHG002x2_19s005136-1-1_Hasenfeld_lane1HG002x02PE20482_2_sequence.txt.gz | NotHighQuality |
| HWVTJAFXY_HG002x1xHG002x2_19s005136-1-1_Hasenfeld_lane1HG002x02PE20489_1_sequence.txt.gz | HWVTJAFXY_HG002x1xHG002x2_19s005136-1-1_Hasenfeld_lane1HG002x02PE20489_2_sequence.txt.gz | NotHighQuality |
| HWVTJAFXY_HG002x1xHG002x2_19s005136-1-1_Hasenfeld_lane1HG002x02PE20493_1_sequence.txt.gz | HWVTJAFXY_HG002x1xHG002x2_19s005136-1-1_Hasenfeld_lane1HG002x02PE20493_2_sequence.txt.gz | NotHighQuality |
| HWVTJAFXY_HG002x1xHG002x2_19s005136-1-1_Hasenfeld_lane1HG002x02PE20495_1_sequence.txt.gz | HWVTJAFXY_HG002x1xHG002x2_19s005136-1-1_Hasenfeld_lane1HG002x02PE20495_2_sequence.txt.gz | NotHighQuality |
| HWVTJAFXY_HG002x1xHG002x2_19s005136-1-1_Hasenfeld_lane1HG002x02PE20496_1_sequence.txt.gz | HWVTJAFXY_HG002x1xHG002x2_19s005136-1-1_Hasenfeld_lane1HG002x02PE20496_2_sequence.txt.gz | NotHighQuality |

### Table S3: Verkko NA24385 Titration Scaffold auN

| % High-Quality Libraries | Repetition | Scaffold auN (Mbp hpc) | Fraction of 192 Library Reference auN (%) |
|---|---|---|---|
| 0 | 1 | 86.9 | 86 |
| 25 | 1 | 94.3 | 93 |
| 25 | 2 | 97.8 | 96 |
| 25 | 3 | 85.2 | 84 |
| 25 | 4 | 94.4 | 93 |
| 50 | 1 | 95.4 | 94 |
| 50 | 2 | 96.8 | 95 |
| 50 | 3 | 99.6 | 98 |
| 50 | 4 | 95.4 | 94 |
| 75 | 1 | 99.9 | 98 |

| | | | |
|---|---|---|---|
| 75 | 2 | 99.6 | 98 |
| 75 | 3 | 97.8 | 96 |
| 75 | 4 | 97.8 | 96 |
| 100 | 1 | 99.9 | 98 |

Table S4: NA24385 hifiasm HiFi Only (44.5x) Phased Assembly Evaluation Statistics

| **Evaluation** | **Metric** | **Graphasing** | **Trio** |
|---|---|---|---|
| Contiguity | auN (Mbp) | 66.0 | 67.9 |
| Contiguity | N50 (Mbp) | 57.8 | 64.9 |
| yak trioeval | Switch Error Rate (%) | 0.23 | 0.16 |
| yak trioeval | Hamming Error Rate (%) | 0.57 | 0.15 |
| yak qv | QV | 56.1 | 55.8 |
| paftools misjoin | Gaps [acrocentric-only] (n) | 7 [0] | 2 [0] |
| paftools misjoin | Interchromsomal Misjoins [acrocentric-only] (n) | 0 [3] | 0 [4] |
| paftools misjoin | Inversions [acrocentric-only] (n) | 1 [0] | 0 [0] |
| paftools asmgene | MMC (%) | 12.0 | 1.4 |
| paftools asmgene | MSC (%) | 1.4 | 0.2 |
